# An impedance-based chemiresistor for the real-time detection of gut microbiota-generated short-chain fatty acids

**DOI:** 10.1101/2022.09.11.507374

**Authors:** Adel Yavarinasab, Stephane Flibotte, Sijie Liu, Carolina Tropini

## Abstract

Short-chain fatty acids (SCFAs) are key molecules produced by gut bacteria in the intestine, that are absorbed into the bloodstream and strongly influence human health. SCFA disruption and imbalances have been linked to many diseases; however, they are seldom used diagnostically as their detection requires extensive sample preparation and expensive equipment. In this work, an electrochemical sensor was developed to enable real-time, quantitative measurement of SCFAs from complex samples in liquid phase without the need for extraction, evaporation, or destruction. An impedance-based sensor for *in vitro* detection of acetic acid, propionic acid, and butyric acid (accounting for more than 95% of SCFAs in the intestine) was fabricated by the deposition of a ZnO and polyvinyl alcohol (PVA) on the surface of a microfabricated interdigitated gold electrode. The sensor was first exposed to a broad, physiologically relevant range of concentrations of SCFAs in isolation (0.5–20 mg/ml) and unlike previously published SCFA sensors that could detect only in gas form with the aid of evaporation, it was able to detect them directly in the liquid phase at room temperature. Electrochemical impedance spectroscopy analysis was then applied to the mixture of SCFAs prepared at different ratios and in complex media at concentrations ranging from 0.5 to 10 mg/ml, which showed the capability of the sensor to measure SCFAs in experimentally relevant mixture. The recorded faradaic responses were then used to train a fit-to-data model to utilize the sensor to screen human bacterial isolates and detect which species secrete SCFAs *in vitro*. This work will allow for the rapid and non-destructive determination of the levels of SCFAs in complex biological samples, providing a miniaturized, highly stable, and highly sensitive sensor for real-time monitoring applications.

## 1 Introduction

The gut microbiota constitutes a large community of microbes that is in constant crosstalk with its mammalian host. Gut bacteria produce many bioactive metabolites which are absorbed by the gut lining and transported into the portal blood system [1,2]. Some of the most important bioactive compounds produced by the gut microbiota are short chain fatty acids (SCFAs), which are mainly produced through the bacterial fermentation of non-digestible carbohydrates in the diet (e.g., plant fiber) and exert significant effects on host metabolism and protection against invading pathogens [3,4]. The most abundant SCFAs in the gut are acetate, propionate, and butyrate, produced by a wide range of bacterial species [5]. SCFAs play a significant role in regulating the integrity of the intestinal tissue, are the main food source of energy for colon tissue cells (colonocytes) and are implicated in regulation of inflammation [6]. In addition to their general role in gut health, SCFAs are absorbed into the bloodstream where they are known to affect multiple organs. They are speculated to mediate crosstalk between the gut and brain (e.g., cognition and emotion) [7], and affect lipid accumulation and obesity by affecting intestinal energy harvesting [8]. Furthermore, increasing evidence supports a regulatory role for SCFAs in skeletal muscle and bone mass [9] and insulin sensitivity [10]. The critical role of SCFAs in human physiology highlights the necessity for accurate detection methods to better understand these prevalent molecules and their role in health and disease.

The conventional method for the detection of SCFAs in biological samples is chromatography. Gas chromatography (GC), first introduced in 1952 for fatty acids [11], is often coupled with a flame ionization detector [12] or a mass spectrometer [13], and uses a free fatty acid phase with a specific capillary column to analyze SCFA acidified water. While this method provides very high sensitivity to SCFA in a solution, it has significant drawbacks such as thermic degradation and consequent structural modification of fatty acids during the methyl esterification process as well as low recovery [14], long incubation time [15], and sample destruction [16]. To address these issues, pre-treatment methods, either physical (e.g., filtration, centrifugation, or dilution) or chemical (acidification with oxalic acid [17], sulfuric acid [18], phosphoric acid [19], etc., or sample distillation [20]), are used; however, such techniques can lead to impurities or column contamination, are time-consuming, and result in loss of column quality over short periods of time [14].

High performance liquid chromatography is the next most common chromatographic method for SCFA detection, which requires derivatization (with 2-nitrophenylhydrazine [21], 3-nitrophenylhydrazine [22], etc.) or deproteinization (with perchloric [23] or sulfuric [24] acid). Other analytical techniques, such as nuclear magnetic resonance [25] and capillary electrophoresis [26], have also been characterized for quantification of SCFAs in biological samples, but present significant issues associated with high cost, low reproducibility and repeatability, and inability of performing real-time analyses due to the long sample preparation time.

Real time analysis of SCFAs would be an important advancement in the medical field by allowing immediate and direct reporting of the availability of these critical compounds. Among the techniques that allow real time detection, electrochemical sensors, and more specifically chemiresistors, provide extremely high selectivity [27]. They also come with other benefits such as ease of fabrication and miniaturization, small footprint, high stability and high temporal-spatial resolution, which are all important for real-time monitoring of biochemical molecules [27]. Chemiresistors are made up of a conductive thin-film sensing layer sandwiched by a pair of electrodes. Upon exposure to the analyte, the electrical properties of the sensing layer change primarily due to resonant quantum tunnelling and give rise to a response. The functionality and efficiency of chemiresistors are heavily dependent on the sensing layer. Carbonaceous materials [28], perovskite-structured (e.g., LaFeO_3_ [29], MgGa_2_O_4_ [30]), and metal oxide (ZnO [31], In_2_O_3_ [32], SnO_2_ [33]) semiconductors (usually doped or with a heterojunction structure) are among the most common class of materials used for SCFA detection. The reaction between the chemisorbed oxygen on the surface of the electrode with SCFAs releases electrons back to the sensing layer, and the resultant decreases in the width of the depletion and accumulation layers change the resistance of the sensor. This has been shown to facilitate SCFA monitoring, i.e., for acetic acid: *CH*_3_*COOH* + 4*O*^-^ → 2*CO*_2_ + 2*H*_2_*O* + 4*e*^-^ or *CH*_3_*COOH* + 4*O*^2-^ → 2*CO*_2_ + 2*H*_2_*O* + 8*e*^-^ [34]. Despite several reports on detection of single SCFAs (e.g., acetate, propionate, and butyrate) in gas phase, there are no reports that differentiate SCFAs in an unaltered biological sample. However, in biological samples different SCFAs are found in liquid phase and within the same sample. Studies on fatty acids in liquid phase are either limited to larger molecules (with greater affinity to bind to the surface) such as indole-3-acetic acid [35] or 2,4-dichlorophenoxy acetic acid [36], or the determination of SCFA has been performed in vinegar and other liquids with pure background [37] and bulky apparatus [38] (Table 1).

**Table 1:** Performance comparison of the developed ZnO/PVA chemiresistor and recent literature for the detection of acetic acid.

| Sensing layer | Phase | RH | Linear Range | Level | Response | T (°C) | Ability to detect AA in a mixture | Biological sample | Ref. |
| --- | --- | --- | --- | --- | --- | --- | --- | --- | --- |
| Y doped ZnO | Gas | - | 1-800 ppm | 1000 ppm | 2.5 | 250 | No | No | [56] |
| Pr-doped ZnO | Gas | dry | - | 800 ppm | 23 | 200 | No | No | [34] |
| Ag-doped ZnO | Gas | - | 1-25 ppm | 100 ppm | 350 | 300 | No | No | [57] |
| Mg-doped ZnO | Gas | - | 20-200 ppm | 200 ppm | 1.32 | 300 | No | No | [58] |
| Y-doped ZnO | Gas | 90 | 1-5 ppm | 1 ppm | 28 | 350 | No | No | [31] |
| SnO2 | Gas | - | 1-100 ppm | 100 ppm | 47 | 260 | No | No | [59] |
| Y-doped ZnO | Gas | - | - | 10 ppm | 50 | 300 | No | No | [60] |
| ZnO/PANi | Gas | - | 1-10 ppm | 10 ppm | 0.22 | RT | No | No | [61] |
| ZnO/PVA | Liquid | 100 | 0.5-20 mg/ml | 0.5 mg/ml | 1.9 | RT | Yes | Yes | This study |

RH= relative humidity; $\text{Response} = \frac{R_{\text{target}}}{R_{\text{initial}}}$ R=impedance measurement at the corresponding concentration; AA=acetic acid; real sample=tested in an actual complex fluid in the presence of living cells

To overcome these limitations, we developed an electrochemical impedance-transduced chemiresistor for determination of SCFAs produced by bacteria directly from biological samples without the need for vaporization or extraction. The sensing layer, made of ZnO/polyvinyl alcohol, was chosen to not have response from other significant by-products of the gut microbiome. To characterize the sensor response with pure SCFAs, the composite was first directly exposed to isolated SCFAs in liquid media. The complexity of the media was then increased, and the sensor was tested in common bacterial growth media with added pure SCFAs. The sensor was then tested in a mixture of SCFAs in a physiologically relevant range of concentrations, and a fit-to-data model was established to predict the Nyquist data of real samples diluted in the electrolyte. Finally, the sensor was used to measure the level of SCFAs secreted by 6 strains of common human bacterial gut isolates grown in the liquid medium: 1 strain of *Bacteroides thetaiotaomicron*, 3 strains of *Lactiplantibacillus plantarum* and 2 strains of *Escherichia coli*. The measurements were validated using mass spectrometry. Measurements with the novel sensor showed that different strains of *L. plantarum* (A138, HA119, and Lp-115) produce the same level of different types of SCFAs; however, this amount is significantly different for strains of *E. coli*. The results were corroborated with mass spectrometry results. Importantly, this paper presents a novel simple method to detect the real time production of important bacterial metabolites directly from complex biological samples providing is a promising new tool for future diagnostics research.

## 2 Experimental section

### 2.1 Sensing layer fabrication

All reagents used were of analytical grade, and were not purified, unless otherwise described. A stock solution of 0.5 g of ZnO nanopowder (<100 nm particle size, Sigma Aldrich, ON, Canada) was prepared in 25 mL ethanol and ultrasonicated with PVA (99+% hydrolyzed, Sigma Aldrich, ON, Canada) for 12 h at different mass ratios (1:5, 1:2, 1:1, 2:1, 5:1). After vortex mixing for 10 mins (Maxi Mix II, Fisher Scientific), the suspension was then deposited dropwise on the interdigitated gold substrate and allowed to calcinate at 80°C on a hotplate for 12 h. The gold electrode, serving as the working electrode, contained two terminals on a glass substrate with a 10 µm gap (0.0188 cm^-1^) and 250 digits (125 digits on each side), each was 6760 µm long. The drop wise method was performed by occupying the least cross-sectional area for hampering the formation of capacitance at low frequencies. The surface morphologies and crystallinity of the ZnO/PVA nanocomposite were investigated with scanning electron microscopy (SEM) and X-ray diffraction (XRD) analysis (**Figure 1**).

**Figure 1.**
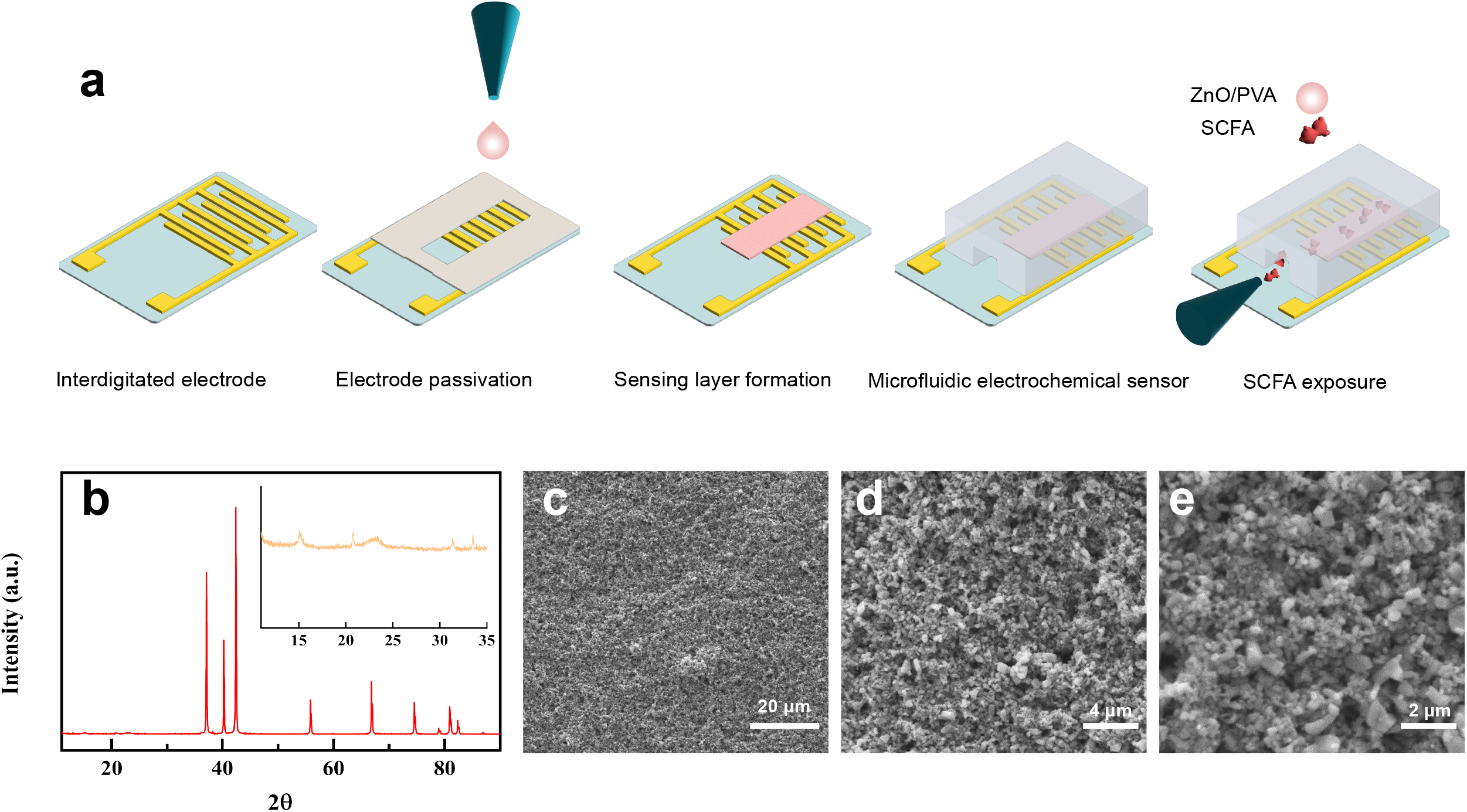
**a**. Schematic view of the microfluidic setup and sensing layer synthesis **b**. X-ray diffraction pattern (inset: 2θ = 12–35° and the effect of PVA on shifting the peaks) **c-e**. Magnified images (SEM) of the sensing layer, showing a homogeneous distribution of ZnO nanoparticles in PVA matrix.

### 2.2 Electrical measurement

The SEM was performed on an FEI Quanta 600 tungsten filament system, operated at 10 kV. The images were collected with the secondary electron detector. The XRD blurb was conducted on a Bruker D8 Advance Bragg-Brentano diffractometer equipped with an Fe filter foil, 0.6 mm (0.3°) divergence slit, incident- and diffracted-beam Soller slits and a LynxEye-XE detector. Continuous-scan X-ray powder-diffraction data were collected over a range 2θ=10-90° with CoKα radiation. The long fine-focus Co X-ray tube was operated at 35 kV and 40 mA, using a take-off angle of 6°. The counting times was performed with a strip detector. The step size was 0.02° with an exposure time at each step of 0.5 s, but in aggregate each step gets a total of 96 s exposure (0.5 times 192 strips).

The Electrochemical measurements were conducted by electrochemical impedance spectroscopy (EIS), using a potentiostat (CH Instruments CHI660D) with a sinusoidal wave with a bias potential of 5 mV over the sweeping frequency range of 100 kHz to 10 Hz with a quiet time of 2 s. The voltage of 5 mV was chosen to minimize the disturbance of the sensing layer and enhance the signal-to-noise ratio. For the experiments conducted in the electrolyte, the solution of 5 mM *K*_3_*Fe*(*CN*)_6_, 5 mM *K*_4_*Fe*(*CN*)_6_, and 0.1 M *KCl* (Sigma Aldrich – Missouri – United States) was used as the redox mediator to transform electrons from the SCFAs to the working electrode. All experiments were conducted at room temperature, with at least 3 replications performed under the same experimental condition (e.g., all at room temperature or pH). After each measurement, the sensor was flushed with deionized water for 1 min to wash away possible residuals from the previous experiment. To prepare mixed SCFA samples, the initial concentration of the butyric acid was set, and the appropriate level of propionic acid and acetic acid was added to the solution and stirred for faster diffusion. For automation and to reduce the amount of analyte required, the measurements of bacterial samples were conducted in a PDMS-based microfluidic chamber which was mounted on top of the gold substrate. Here, the mold was fabricated on a silicon wafer using standard photolithography containing of a straight chamber, one inlet and one outlet with L 10.0 × W 0.5 × H 0.5 mm dimensions. The SYLGARD 184 silicone elastomer base and curing agent (Dow Corning Co. Ltd) with a ratio of 1:9.8 was poured onto the mold. The PDMS was then peeled off and plasma-treated for 1 min at 50 W power with the interdigitated electrode, followed by heating overnight at 60°C to achieve bonding between the microfluidic chamber and the sensor with no leakage. The schematic view of sensing platform is shown in **Figure 1a**.

### 2.3 Cell culture

#### Culturing conditions and isolation of bacteria from probiotics

Lysogeny Broth and Mega Medium (LB and MM, Table S1) were chosen as media types that support the growth of diverse species of bacteria. MM has SCFA in the base medium that support the growth of specific fastidious bacteria [39]. *E. coli* strains were streaked onto Luria Broth (LB Fisher BP1426500-500) + 1.5% agar (Fisher BP1423500) plates and grown overnight at 37ºC. Single colonies were inoculated into 5 mL liquid LB and grown overnight at 37ºC shaking at 225 rpm. *B. thetaiotaomicron VPI-5482* (*B. theta*, ATCC 29148; GenBank AE015928.1) was grown in an anaerobic chamber (gas supplied by Linde: NI CD5H11U-T [5% CO_2_, 5% H_2_, 90% N_2_]). All plates and media used to grow *B. thetaiotaomicron* were pre-reduced under anaerobic conditions for at least 12 hours before use. *B. thetaiotaomicron* was streaked onto BHIS-agar plates (Brain-Heart Infusion broth [BD B11059] supplemented with 0.5 mg/mL vitamin K1 (Alpha Aesar AAL1057506) and 5 μg/mL hemin chloride (MilliporeSigma 37415GM, with 1.5% agar), and grown at 37°C overnight. Single colonies were inoculated into 5 mL liquid BHIS broth and grown at 37°C overnight without shaking. Liquid overnight cultures were inoculated into 5 mL MM. *Lactobacillus*-containing probiotic pills (Super 8 Plus Probiotic, FLORA) and (Gentle-Care Probiotic, Genuine Health) were purchased from local pharmacies. The probiotics were in the forms of capsules, pellets, and powder, and they were brought into the same anaerobic chamber used for *B. thetaiotaomicron* culturing. The probiotics were placed in individual 15 ml centrifuge tubes and dissolved in pre-reduced, sterile phosphate-buffered saline (PBS). The dissolved probiotics were struck on De Man, Rogosa and Sharpe (MRS) agar and incubated at 37ºC for 24 hours [40]. The colonies were resuspended in liquid media for DNA extraction to verify the identity of the bacteria by 16S rRNA sequencing [41].

#### DNA Extraction from Colonies and 16S rRNA Sequencing

Bacterial DNA was extracted using a DNeasy Blood and Tissue Kit (Qiagen, ON, Canada) following manufacturer instructions. Amplicons of the 16S ribosomal RNA (rRNA) coding sequence region were generated using prokaryotic primers broad-range bacterial primer 8F (5’-AGA GTT TGA TCC TGG CTC AG-3’) and universal primer 1391R (5’-GACGGGCGGTGTGTRCA-3’) using a standard PCR protocol [42]. Purified PCR products were sequenced at the Sequencing + Bioinformatics Consortium at the University of British Columbia. Blast sequence analysis tool as used to align the resulting sequences against the NCBI database to identify the specific isolated strain [43].

### 2.4 Fit-to-data model

Impedance data obtained with mixtures of analytes at various concentrations were explored and visualized in R [R: A language and environment for statistical computing. R Foundation for Statistical Computing, Vienna, Austria. URL https://www.R-project.org/]. Multiple linear regression analyses were performed with the function *lm* and the results plotted with the help of the packages *ggeffects* [44] and *ggplot2* [45]. The metrics quantifying the quality of the fit for the mixture experiments (R^2^ and p-values) were part of the output of the fitting function *lm* and they were accessed using the summary function (https://search.r-project.org/R/refmans/stats/html/summary.lm.html) in R. Code can be found at https://github.com/Tropini-lab/Adel-s-SCFA-sensor.

### 2.5 Gas chromatography tandem mass spectrometry

Agilent 8890 gas chromatograph coupled with an Agilent 7010B triple quadrupole mass spectrometer with CTC PAL autosampler equipped with headspace and SPME options and a split/splitless injector was utilized for the gas chromatography. The column used was an Agilent DB FATWAX UI 30-meter column, 0.25 mm diameter and 0.25 μm film thickness with helium as carrier gas at a flow of 1.2 ml/min. Samples and standard curves of underivatized SCFA were analyzed by headspace injection after incubation in acidified water at 95°C for 40 minutes and utilized a 0.5 ml injection and a 10:1 split. The column oven was temperature programmed from 90°C to 230°C over 18 minutes with baseline separation of all SCFAs. Calibration curves utilized authentic standards and deuterium labelled internal standards of acetic, propionic, butyric and caproic, all purchased from Millipore Sigma or CDN isotopes. The GCMS was operated in SRM mode and regression lines were calculated using quadratic fits with correlation coefficients of 0.995 to 0.9995.

## 3 Results and discussion

### 3.1 Film characterization

To examine the crystal structure and crystallinity of the ZnO/PVA sensing layer, XRD measurements were performed (**Figure 1b**). The polycrystallization with hexagonal wurtzite (diamond) structure of ZnO was well-indexed by three characteristic sharp peaks at 2θ = 38°, 40°, and 42°, attributed to the (100), (002) and (101) crystal planes, respectively. Reflection peaks related to ZnO at 2θ = 56°, 67°, 75°, 79°, 82°, and 84° correspond to the reflections from the next crystal planes: (102), (110), (103), (112), (201), and (202), respectively, in agreement with available literature values (JCPDS card no. 36-1451). PVA, however, is a partially crystalline polymer due to the intermolecular hydrogen bonding within the network. The small hump around 22° corresponds to the (101) plane of PVA, signifying its semi-crystalline nature. The ZnO content suppresses the crystallinity of PVA and only a broad depressed peak was observed at the (101) plane at 2θ = 20 – 25°. This was attributed to the weakening of hydrogen bonding in the PVA chain and the reduction of the dense structure of molecular packing [46]. The overall XRD pattern of the PVA/ZnO composite revealed a slight shift (4°) of all peaks (2θ) due to the incorporation of PVA in the ZnO crystal lattice and using a cobalt source instead of the commonly used copper for the measurement. There were no peaks related to impurities or mixed compounds, supporting the effective incorporation of the ZnO lattice in the PVA matrix. The formation of a dispersed composite structure due to the difference between the crystal structure of ZnO and PVA is a prominent parameter for the sensing characteristic of the sensor.

To study the surface morphology of the sensing layer, SEM was performed (**Figure 1c-e**). SEM images of ZnO (white nanoparticles) embedded in PVA matrix (darker background) demonstrated a fairly uniform and homogeneous distribution of ZnO (mean diameter of about 50 nm) throughout the polymeric matrix: uniform pores and high surface area were observed throughout the surface. ZnO nanospheres tended to agglomerate due to their high surface area, particularly for smaller nanoparticles, exhibiting a specific ZnO/PVA structure. Of note, while different mass ratios of the metal oxide:polymer were tested (1:5, 1:2, 1:1, 2:1, 5:1), the percolation threshold was achieved at 1:1, and thus all experiments were conducted at this ratio in this paper.

### 3.2 Impedance spectroscopy of individual SCFAs

The functionality of the developed ZnO-reinforced PVA-based sensor was first analyzed by the introduction of the SCFA acetic acid dissolved in the electrolyte at a broad range of 0.5 to 20 mg/ml (∼ 8 mM – 333 mM). The range of the concentration was chosen to span the acetate levels in healthy people, patients suffering from fatty liver and steatohepatitis [47], as well as colitis-induced mice [48]. **Figure 2** shows to the Nyquist spectrum of the chemiresistor, which forms a quarter circle connecting with a strike line at low frequencies. As the radius of the Nyquist is attributed to the charge transfer resistance in alternative current (AC) (compared with the resistance in direct current [DC] models), one can infer that an increase in the concentration of acetic acid, and localization in the surface pores, results in higher conductivity of the chemiresistor. Specifically, as an n-type semiconductor, the adsorbed oxygens trap electrons from the conduction band of ZnO and form a depletion region. The chemisorbed oxygens oxidize the acetic acid, and the trapped electrons move to the conduction band, decreasing the resistance of the film [34]. While the band theory is operational at elevated temperatures (due to the dependence of reaction kinetics on temperature), the integration of PVA particles (with a significant contribution to the composite) forms a heterojunction with a hole-electron depletion layer at the interface of ZnO and PVA [49]. As such, the migration of free electrons from ZnO toward PVA and diffusion of the holes of PVA to ZnO contributes a new balance between the energy bands and Fermi level. This provides a higher surface area for the sensing layer to interact with acetic acid and facilitates molecular diffusion at the grain boundaries at room temperature [49]. The calibration curve of the ZnO-PVA chemiresistor upon exposure to acetic acid at a frequency of 1000 Hz is represented in **Figure 2d**, showing a linear response, *Z*_*AA*_ = -2.303*C*_*AA*_ [*mg*/*ml*] + 178.4, where *Z*_*AA*_ is the magnitude of impedance and *C*_AA_ is the concentration of dissolved acetic acid in the electrolyte in mg/ml. The small error bars (standard deviation from the mean values) and high R^2^ value (∼0.97, *p* - *value <* 0.0001) demonstrate the reliability of the measurements.

**Figure 2.**
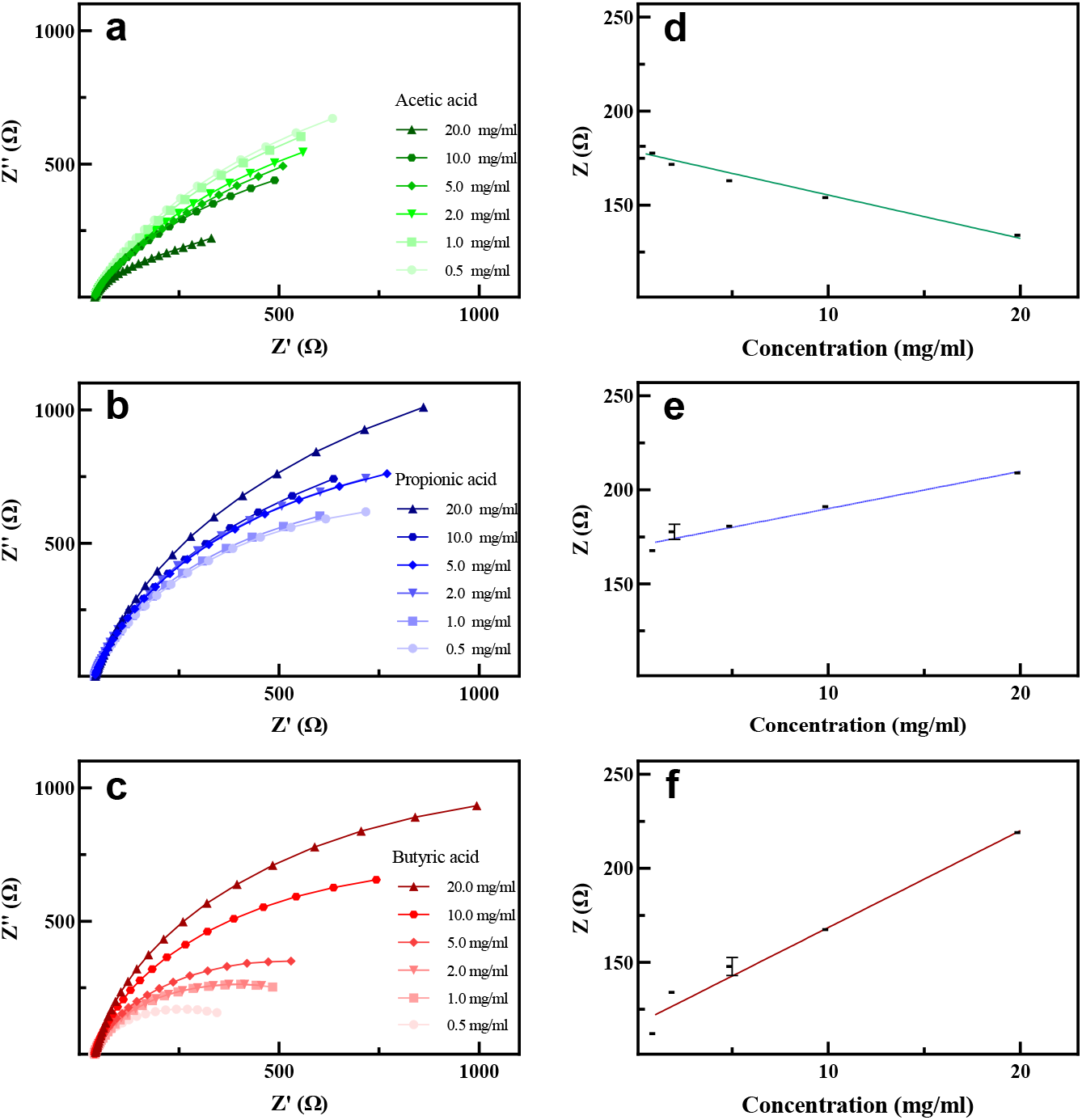
The Nyquist spectra of **a**. acetic acid, **b**. propionic acid, and **c**. butyric acid **d-f**. the corresponding calibration curve (the magnitude of impedance) at a frequency of 1000 Hz at SCFA concentrations of 0.5–20 mg/ml.

Having demonstrated sensitivity to acetic acid, the sensor was then exposed to two other medically relevant SCFAs: propionic acid, and butyric acid in physiological ranges of 0.5 to 20 mg/ml (∼ 20 – 270 mM) [47,48]. The real and imaginary components of the impedance of the sensing layer after exposure to butyric acid and propionic acid at varying concentrations follow the same trend as for acetic acid; however, increasing the concentration of propionic and butyric acids increases the charge transfer resistance of the sensor and consequently the radius of the Nyquist plot (**Figure 2b, c**, and **Figure 2e, f**). We interpret this result to indicate that due to the small dipole moment between the C-H bond and the larger nonpolar parts of propionic acid and butyric acid, higher concentrations of such SCFAs lead to increases in the resistance of the sensing layer. Importantly, the film still elucidated a linear response to the analytes of interest with calibration curve equations of *Z*_PA_ = 1.985*C*_*PA*_[*mg*/ *ml*] + 170.1 (*R*^2^ = 0.95, *p* - *value* < 0.0001) and *Z*_*BA*_ = 5.157*C*_BA_ [*mg*/*ml*] + 116.8 (*R*^2^ = 0.97, *p* - *value* < 0.0001), for propionic acid and butyric acid, respectively.

### 3.3 Reproducibility, stability, and selectivity of the chemiresistor-on-a-chip

To study the selectivity of the ZnO-PVA chemiresistor, the sensor was introduced to 5 mg/ml of molecularly similar-structured compounds, ethanol, ethanol, and isopropanol (IPA). These chemicals are found at low level in the gut [50,51]. As illustrated in **Figure 3a**, the Nyquist plot formed the classic pattern of a depressed half circle at high frequencies followed by a strike line as a common faradaic probe for the analytes [52]. In short, the half circle is modelled by the double layer capacitance and/or constant phase element (formed by ion solvation) as well as the Schottky contact/charge transfer resistance, while the straight line is attributed to diffusion (Warburg impedance) [53]. The impedance-frequency characteristics of the sensing layer upon exposure to SCFAs and alcohols show a notably different behaviour, suggesting the selectivity of the sensor for the analytes of interest.

**Figure 3.**
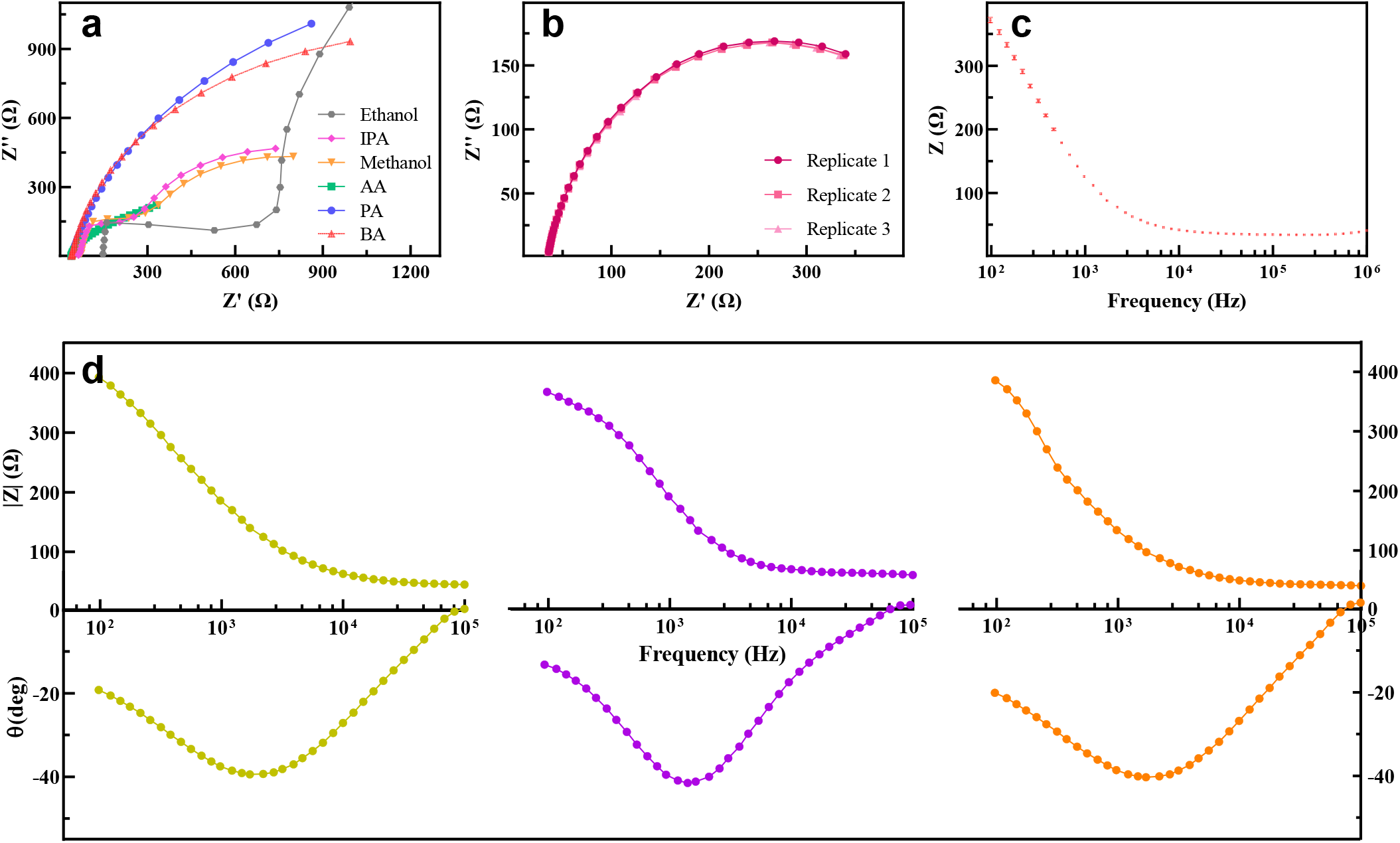
**a**. The different Nyquist spectra of the SCFAs (IPA= isopropanol, BA = butyric acid, AA = acetic acid, PA = propionic acid) and other analytes of interest at 20 mg/ml suggest high selectivity of the chemiresistor **b**. Relative change in the Nyquist spectra and **c**. the Bode plot (impedance vs. frequency) of the chemiresistor after 3 consecutive exposures to 0.5 mg/ml butyric acid, suggesting recovery of the sensor. The error bars in **c** show standard deviation. **d**. The similar Bode spectra (impedance and phase angle vs frequency) of the sensor exposed to 0.5 mg/ml butyric acid with 3 simultaneously fabricated sensors demonstrates the reproducibility of the sensor at low SCFA concentrations.

The repeatability of the results and recovery of the sensor was investigated through the response of the thin film toward 0.5 mg/ml butyric acid after 3 consecutive measurements in the electrolyte under the same experimental conditions, and no significant deviation was observed (**Figure 3b, c**). The standard deviation and relative standard deviation of impedance at a frequency of 1000 Hz were 0.085 Ω and 0.0020 Ω, respectively, showing repeatability and the ability of the thin film to recover to its initial condition and produce comparable results.

After assessing repeatability, we investigated the reproducibility of the sensor by synthesizing the sensing layer 3 times and exposing each sensor to 0.5 mg/ml butyric acid dissolved in the electrolyte. The Bode (impedance and phase angle vs frequency) and Nyquist plots of the 3 sensors are represented in **Figure 3d** and **Figure S1** in the frequency range of 10^5^ to 10^2^ Hz. Of note, the formation of additional stray capacitance at frequencies higher than 10^5^ and the following discrepancy may result in a positive phase angle while no inductance appears to be generated (data not shown here), and therefore data collected only using frequencies below 10^5^ were only analyzed [52]. The maximum value of phase angle (40°) occurred in the frequency range of 10^3^ to 10^4^ Hz, suggesting the frequency dispersion and the interfacial heterogeneity of the sensing layer [54]. The value of phase angle suggests that the capacitive behaviour of the sensing layer acts as an open-circuit and equals the source voltage [55]. In addition, the magnitude of the impedance gradually increased, as a common impedance response in faradaic probes. The negligible deviation (the average impedance of 392.66 Ω at the frequency of 1000 Hz with the relative standard deviation of 0.016 Ω) between the 3 separately developed sensors following exposure to butyric acid confirmed the high reproducibility of the method. The sensing performance of the fabricated sensor was calculated through the response of the sensor to 20 mg/ml propionic acid over the base response 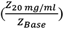 within a period of 16 days (**Figure S2**). The sensor showed a stable response with an average response, standard deviation, and relative standard deviation of 1.92, 0.03, and 0.017, respectively, at a frequency of 1000 Hz (**Figure S2**). The base response equal to 1 is defined as the sensor exposed just to the electrolyte.

The developed sensing system outperforms Recent ZnO-based SCFA sensors as it can directly detect the analyte in the liquid phase and at room temperature with similar sensitivity which are required for medical applications (Table 1). The high performance of the sensor is attributed to the highly porous film morphology of the composite, which takes advantage of facilitated electron transfer via the 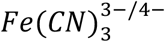 and *KCl* electrolytes. In comparison, the traditional metal oxide chemiresistors need sample evaporation at elevated temperatures, which signifies energy waste and might be influenced by long-term drift issues due to the sintering effect.

Furthermore, the developed sensor can directly detect the level of SCFAs in real aqueous samples with only a small sample requirement by confining the measurement inside a microfluidic device such that precious and significantly low-volumes samples can be tested. Specifically, while regular GC analysis required at least 0.5 ml of samples in liquid medium, the total volume of the chamber was around 0.0025 ml, or a 200-fold volume reduction. Indeed, smaller chambers (with the cross-sectional area of W 50 × H 50 μm) could also be fabricated to further reduce use of samples for specific applications.

### 3.4 Impedance spectroscopy of SCFAs in bacterial media

Having established the sensor reproducibility and stability, more complex samples were analyzed using complex bacterial growth media added directly to the electrolyte base. For determination of SCFAs from different bacterial strains, measurements were conducted in LB – a common bacterial culture medium for *E. coli* [62], and MM – a rich medium known to support the growth of diverse strains [39] (Experimental section). As both LB and MM inherit weak electron transfer characteristics, different dilutions of medium and electrolyte were prepared and tested and a volumetric ratio of 30:1 (electrolyte:medium) was chosen for further experiments (**Figure S3**). Of note, the starting point of the Nyquist diagram represents the bulk resistance of solution, and higher dilution with LB and MM increased the resistance of the solution and decreased the conductivity of the film. This is due to the lower levels of salts used in growth media to support bacterial growth, compared to the electrolyte solution. SCFAs dissolved in the electrolyte/LB or MM were then measured with the chemiresistor, and Nyquist spectra were recorded (**Figure 4**). The sensor followed a similar trend as for SCFAs dissolved in just electrolytes (**Figure 2**). One major difference is that no significant response was observed for butyric acid in LB (**Figure S4**), which can be attributed to the weak conductive characteristic of LB and the nonpolar nature of butyric acid, hindering electron transfer from butyric acid to the composite at room temperature and consequently leading to no response. In contrast, the significant concentration of salts found in MM (Table S1) facilitates electron tunnel transfer, and the impedimetric results in MM appeared to be comparable with pure electrolyte, indicating it is a preferable medium compared to LB for obtaining these measurements. Notably, an initial amount of SCFA was already present in MM (Table S1), and the base response was considered on the exposure of the sensor to MM with the dissolved SCFA in it. The more complete curved half circle in MM was attributed to the additional charge transfer in parallel to the double layer capacitance (a common practice in the equivalent circuit of aqueous media [63]) which was generated in the sample because of the significant level of ionic compounds in the background. The linear response of the sensor toward SCFAs in MM at a frequency of 1000 Hz were *Z*_*AA*_ = -49.39*C*_*AA*_ [*mg*/*ml*] + 874.8 (*R*^2^ = 0.96, *p* - *value <* 0.0001), *Z*_*PA*_ = 36.53*C*_*PA*_ [*mg*/*ml*] + 113.4 (*R*^2^ = 0.98, *p* - *value <* 0.0001), and *Z*_BA_ = 37.92*C*_*BA*_ [*mg*/*ml*] + 344.0 (*R*^2^ = 0.94, *p* - *value* < 0.0001) for acetic acid, propionic acid, and butyric acid, respectively. The concentration was based on the added SCFA to MM. The impedance spectra of acetic acid and propionic acid at the concentration of 1.0 to 10 mg/ml in LB are shown in **Figure S5**. Given our results, measurement using the described sensor are most reliable in more conductive media. The linearity of the impedance response within this range is represented **Figure S5c-d** for acetic acid and propionic acid: *Z*_*AA*_ = -2.723*C*_*AA*_ [*mg*/*ml*] + 198 (*R*^2^ = 0.93, *p* - *value* < 0.0001) and *Z*_AA_ = 5.397*C*_AA_ [*mg*/*ml*] + 258 (*R*^2^ = 0.96, *p* - *value* < 0.0001), respectively.

**Figure 4.**
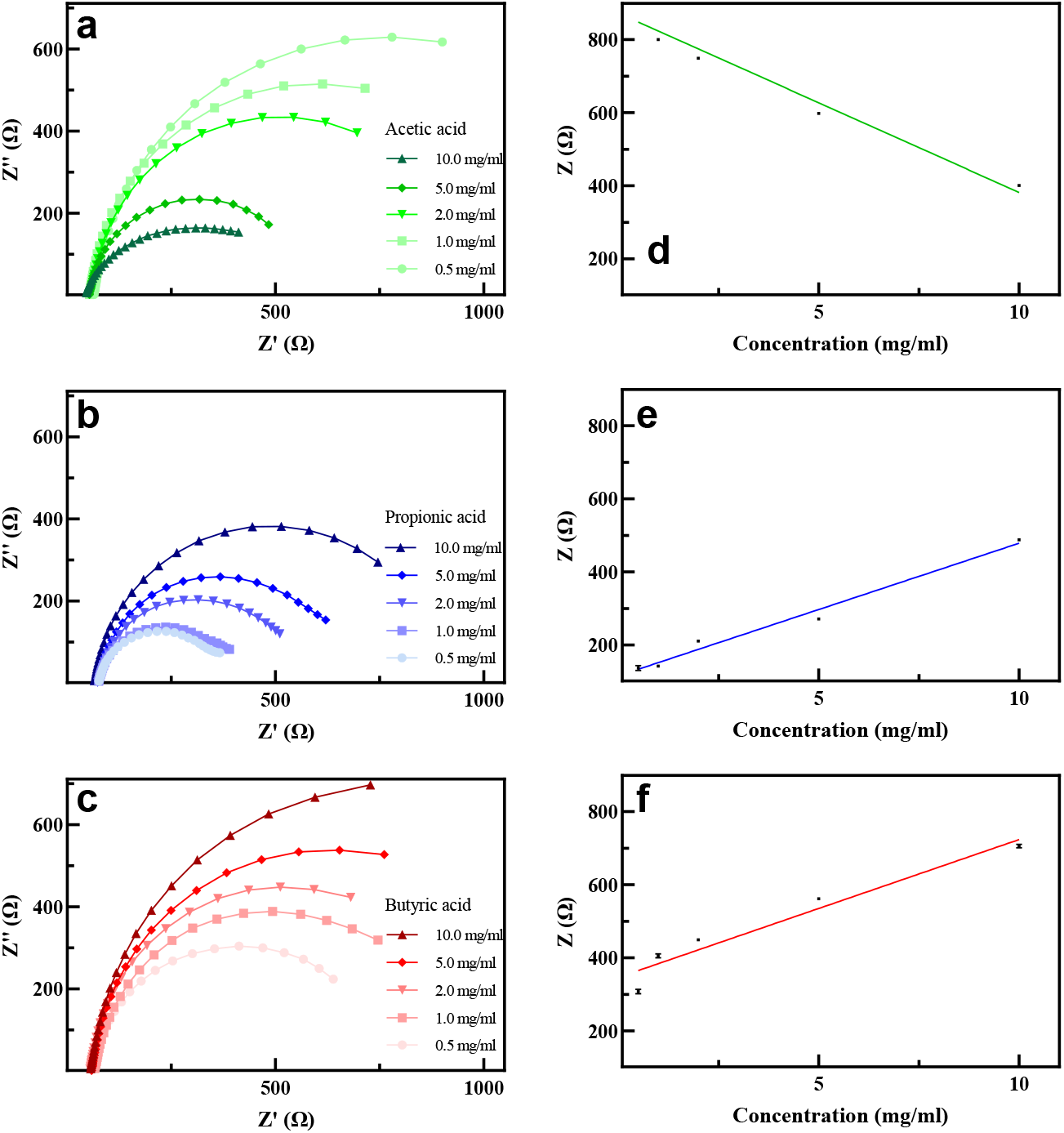
The Nyquist spectra of **a**. acetic acid, **b**. propionic acid, and **c**. butyric acid, dissolved in MM at a volumetric electrolyte:medium ratio of 30:1 and concentration of 0.5–10 mg/ml, and **d-f**. the corresponding calibration curve, follows the same trend as pure electrolyte.

### 3.5 Impedance spectroscopy of SCFAs in mixture and fit-to-data model

Bacteria produce different amounts of SCFAs depending on the species, growth state and condition, and the determination of analytes in a mixture is relevant for practical applications. Therefore, 40 samples at different physiologically relevant SCFA concentrations (in the range of 0 to 10 mg/ml, targeting levels observed in *in vivo* studies [48]) were prepared and the impedance behaviour of the 3 analytes of interest was recorded by changing the concentration of one analyte and keeping the other two SCFAs constant. The concentrations of acetic acid and propionic acid were in the range of 0 to 10 mg/ml while two concentrations were chosen for butyric acid, 0.5 and 1.0 mg/ml (**Figure S6**), as the level of butyric acid is significantly smaller than the other two in biological samples (the molar ratio is of 60:25:15 acetic:propionic:butyric acid in the colon [64]). **Figure 5a** shows representative Nyquist spectrum with the level of butyric acid kept constant at 0.5 mg/ml. While all individual measurements were conducted at 1000 Hz, it was observed that in the SCFA mixture more robust results could be obtained at the frequency of 100 Hz. The imaginary component of impedance (*Z*^”^) at 100 Hz is reported instead of *Z*^’^ as the response is directly associated with the charge transfer resistance [27] and *Z*” maintained a linear relationship with different acid concentrations, indicating its utility in converting the sensor measurements to SCFA concentrations in mixture conditions. For example, the descending value of *Z*^”^ (**Figure S7a-b**) for the impedance changes of acetic acid at the concentration of 10 mg/ml and 0.5 mg/ml of propionic acid and butyric acid and the increasing value of *Z*^”^for propionic acid in the mixture (**Figure S7c-d**) revealed the same trend as it was observed in the MM and pure electrolyte. Results from a multiple linear regression for the experiments with a butyric acid concentration of 0.5 mg/ml are shown in **Figure 5b-c** where the fit *Z*” (at 100 Hz) = 279.850 Ω – (23.607 × acetic acid concentration [mg/ml]) + (40.716 × propionic acid concentration [mg/ml]) achieved a correlation coefficient R^2^ of 0.99 (adjusted R^2^ of 0.9892, p-value < 2.2e-16). To investigate reproducibility, we repeated the experiment with concentrations 5.5-5.0-0.5 mg/ml (acetic-propionic-butyric acid) and obtained the experimental value *Z*” = 358.6 Ω at frequency of 100 Hz and compared with the predicted value according to the fit (353.4 Ω), which corresponding to a difference of only 1.5%. This data shows that the developed chemiresistor can preserve its linearity in an SCFA mixture, and it is sensitive to individual SCFAs as low as 0.5 mg/ml. Our results therefore indicate this sensor can be utilized to detect SCFAs in complex samples as those found in medical applications. Importantly, precise quantifications of acetate, propionate and butyrate in these mixtures will require knowledge of the levels of one of the three SCFA. It should be noted that in biological systems, many microorganisms produce only one or two SCFA significantly. For example, while *Ruminococcus* and *Faecalibacterium* genera are significant producers of butyrate, other probiotics (e.g., *Lactobacillus* genera) strains are usually associated with the secretion of propionate [65]. We therefore envision the use of our sensor as a fast detection system in clinical settings to identify samples that should undergo further analysis.

**Figure 5.**
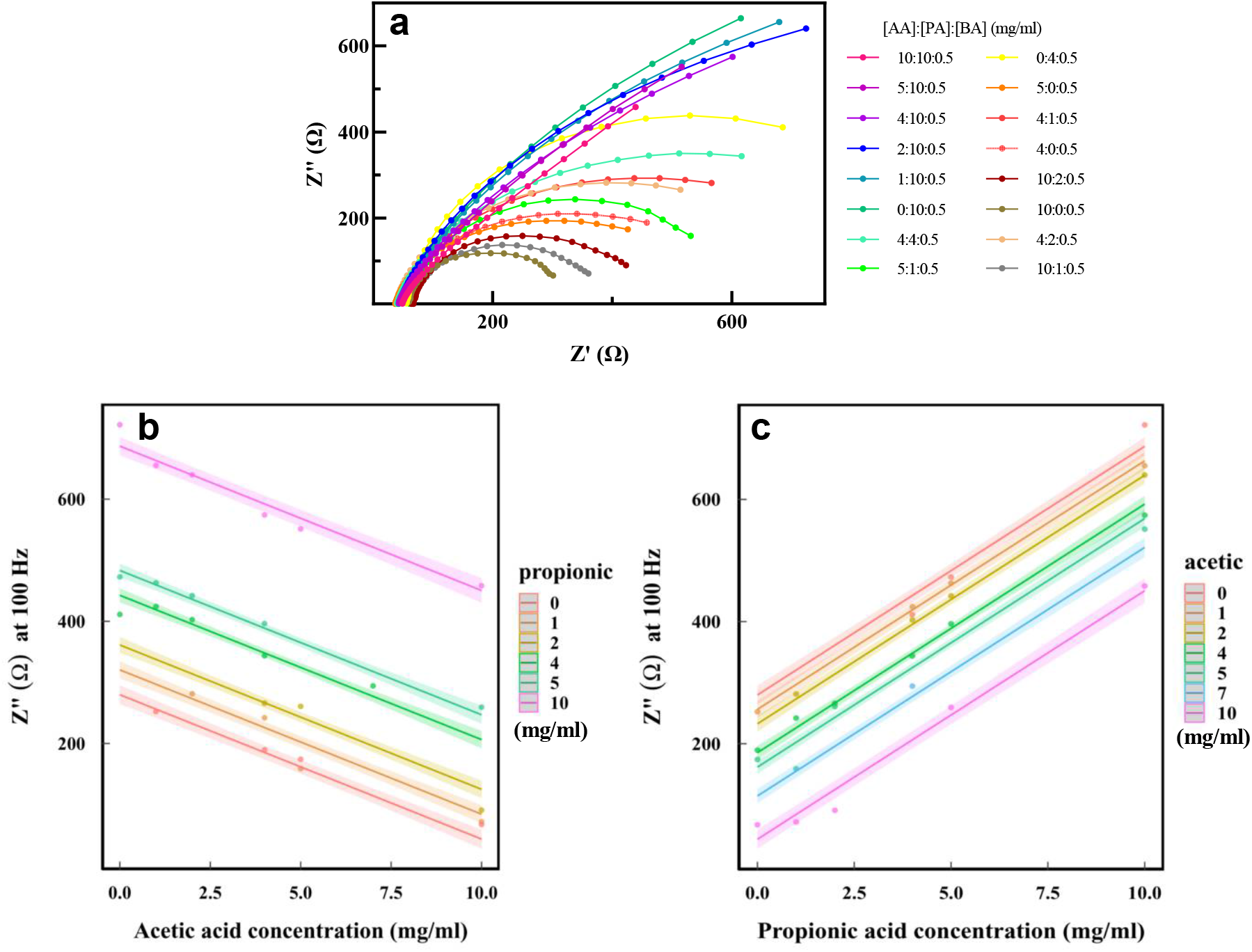
**a**. The Nyquist spectra of the sensing layer at various concentrations of acetic acid and propionic acid in the mixture at the concentration of 0.5 mg/ml butyric acid. The three acid concentrations are listed in the legend (acetic-propionic-butyric). Two dimensional representations of the multiple regression fit to the experiments for **b**. acetic acid and **c**. propionic acid with a butyric acid concentration of 0.5 mg/ml for Z” at the frequency of 100 Hz, R^2^ = 0.99. The ribbons correspond to confidence intervals of 95%.

### 3.6 Live bacterial sample analysis

To analyze the performance of the sensor with previously uncharacterized complex bacterial samples, 6 bacterial strains from three species were grown in their corresponding media (Experimental section). After dilution with the standard electrolyte as above, they were exposed to the electrochemical sensor (**Figure 6a**). The level of SCFAs in 3 strains of *L. plantarum A138, HA119*, and *Lp-115*, 2 strains of *E. coli S17-1 λpir* and *One Shot™ PIR1*, and *B. thetaiotaomicron* (as a control) were measured with the sensor and the values very cross-validated with gas chromatography (GC) after extraction. In *E. coli* strains we expected only one SCFA to be produced in the sample (primarily acetic acid [66,67]), therefore the predicted concentration was achieved by the calibration curve of acetic acid and propionic acid in LB in Section 3.4 at the frequency of 1000 Hz with the confidence interval of 0.99. **Figure 6a** demonstrates that various strains of *E coli* secrete SCFAs at significantly different levels. The small deviation (∼ 0.1 mg/ml for *One Shot™ PIR1* and ∼ 0.03 mg/ml for *S17-1 λpir*) between the response of the sensor and GC can be attributed to other fatty acids (e.g., valeric acid and isovaleric acid) in the background that were produced in insignificant amounts of < 14 μg/ml. The Nyquist spectra of 3 strains of *L. plantarum* showed an extremely similar response (**Figure 6b**), suggesting that the level of produced SCFAs should be very similar. With the obtained value of *Z*” at the frequency of f = 100 Hz and the fit-to-data model, for the fixed concentration of 0.5 mg/ml of butyric acid (as expected from previous studies (e.g., [68]), 3 strains of *L. plantarum* secreted ∼ 2.1 mg/ml of acetic acid and 0.6 mg/ml of propionic acid, which was confirmed by GC. The deviation in the concentration of butyric acid in *L. plantarum* strains is attributed to the generation of isobutyric acid and isovaleric acid at the level of < 0.1 mg/ml. No significant concentration was obtained *B. thetaiotaomicron*, suggesting the negligible presence of SCFAs present in the sample which was proven by GC. These results show that the application of the sensor can be generalized to a broad range of bacteria producing at least 200 µg/ml of total SCFAs.

**Figure 6.**
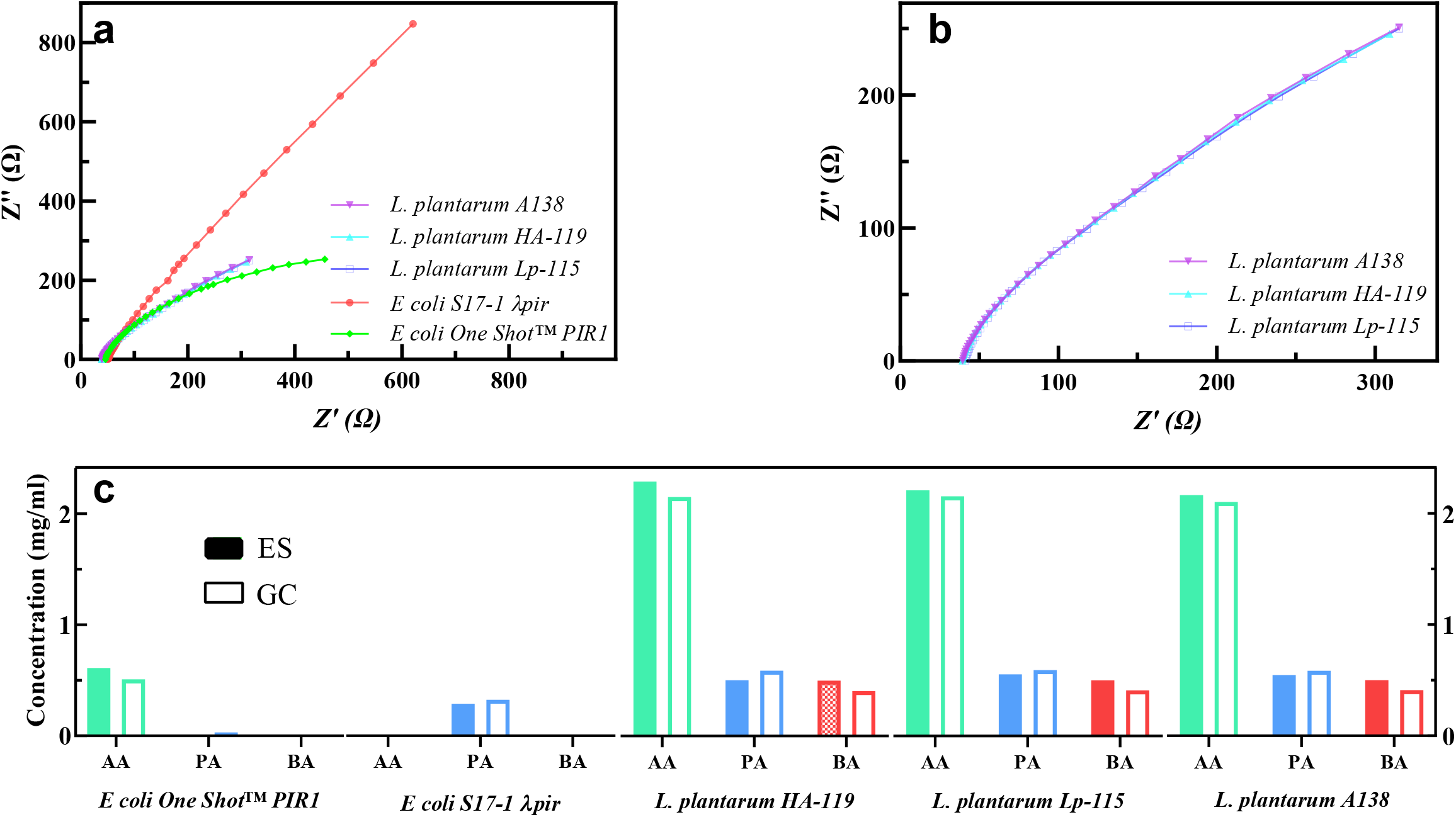
**a**. The Nyquist spectra of 5 strains of *L. plantarum* and E. *coli* in their corresponding medium (AA=acetic acid, PA=propionic acid, and BA=butyric acid) **b**. Similar response between 3 strains of *L. plantarum* **c**. comparison between the determination of the level of SCFA with electrochemical sensor (ES) and gas chromatography (GC) for 5 strains.

## 4 Conclusion

In this study, an impedance-based electrochemical sensor was developed to screen complex human bacterial isolates and detect the individual and total amount of SCFAs secreted from bacterial species *in vitro*. The sensor was synthesized through the formation of a heterojunction between ZnO and PVA. Using EIS, SCFAs in isolation, dissolved in the electrolyte, MM, and LB were introduced to the sensor, and linear responses were measured. To detect the ability of the sensor to determine SCFAs *in vitro*, the sensor was exposed to a mixture of SCFAs over a broad range of concentrations, and a fit-to-data model was used to train a platform from the data obtained for the mixture results. The sensor could detect the level of SCFAs in bacterial isolates (*L. plantarum* and *E. coli)* and identify them with high accuracy with only 2.5 μL of sample required (200 times less than a chromatography method). This technique outperforms the current gold standard method (gas chromatography) as the detection is directly in liquid phase at room temperature without the need for evaporation, extraction, or sample destruction. Importantly, the method is also real-time, performing measurements for one sample in less than 2 minutes. Future applications will require testing of the sensor using more complex human biological fluid samples such as blood where SCFAs may have a biomarker role in multiple disorders [69]. Thanks to its small size, the developed sensor could be used to test *in vivo* samples inside animals where bulky chromatographic devices are normally unable to execute. This method also presents limitations. Specifically, in the case of the presence of other analytes in a significant amount, the signal-to-noise ratio may decrease significantly affecting the performance of the system. In addition, if multiple sources of SCFA were to be present in the biological fluid (such as ones coming from intestinal samples where hundreds of bacterial species are growing), a more comprehensive calibration and predicted model will be required. Finally, in future applications, specific media should be tested to ensure the largest signal possible can be obtained. All in all the method presented here demonstrates that electrochemical sensors can be employed in the real-time measurements of complex biological molecules such as SCFAS without the need of extraction and should be further investigated as the medical relevance of these compounds increases the needs for fast and reliable measurements to be performed at the time of collection.

## Supporting information

Graphical abstract

## Acknowledgments

The authors acknowledge that the land we performed this research on is the traditional, ancestral, and unceded territory of the xwməθkwəy?əm (Musqueam) Nation. The land it is situated on has always been a place of learning for the Musqueam people, who for millennia have passed on in their culture, history, and traditions from one generation to the next on this site. We encourage others to learn more about the native lands in which they live and work at https://native-land.ca/

The authors acknowledge support for this project from CIHR Team Grant: Canadian Microbiome Initiative 2 (FASF19-04421), Crohn’s and Colitis Canada (625155), CIFAR (GS20-011), Michael Smith Foundation for Health Research Scholar Award (18239), Johnson & Johnson Women in STEM2D Award (15007) and Canada Foundation for Innovation (38277). We thank Dr. Nina Maeshima, Giselle McCallum, Dr. Katharine Ng and Dr. Boris Stoeber for critical feedback on the manuscript, Dr. Camden Hunt and Dr. Curtis P. Berlinguette for electrochemical potentiostat station, and Jacob Kabel for surface characterization, SEM, and XRD imaging.

## Figure Legends

**Figure S1** Reproducibility of the sensor through fabrication of 3 chemiresistors exposed to 0.5 mg/ml butyric acid and similar Nyquist spectra.

**Figure S2** The small deviation between the response of the sensor after 16 days of exposure to 20 mg/ml propionic acid.

**Figure S3** The Nyquist spectra of 10 mg/ml acetic acid in **a**. the electrolyte, and electrolyte:LB at a volumetric ratio of **b**. 1:1 and **c**. 30:1.

**Figure S4** The Nyquist spectra of butyric acid dissolved in LB with a volumetric electrolyte:medium ratio of 30:1.

**Figure S5** The Nyquist spectra of **a**. acetic acid, **b**. propionic acid dissolved in LB with a volumetric electrolyte:medium ratio of 30:1, and **c-d**. the corresponding calibration curve. No significant response was observed with butyric acid.

**Figure S6 a**. The Nyquist spectra of butyric acid at two different concentrations of 0.5 and 1.0 mg/ml at various level of acetic acid and propionic acid **b**. The calibration curve of butyric acid at two concentrations represents the non-linear characteristic (different slope) for various level of acetic acid and propionic acid.

**Figure S7 a**. The Nyquist spectra of acetic acid in the mixture of 10 mg/ml propionic acid and 0.5 mg/ml butyric acid and **b**. the calibration curve for *Z* ^”^ at the frequency of 100 Hz **c**. The Nyquist spectra of propionic acid in the mixture of 1 mg/ml acetic acid and 0.5 mg/ml butyric acid and **d**. the calibration curve achieved by *Z* ^”^ at the frequency of 100 Hz.

## References

[1] G.P. Donaldson, S.M. Lee, S.K. Mazmanian, Gut biogeography of the bacterial microbiota, Nat Rev Microbiol. 14 (2016) 20–32. https://doi.org/10.1038/nrmicro3552.

[2] C. Tropini, K.A. Earle, K.C. Huang, J.L. Sonnenburg, The Gut Microbiome: Connecting Spatial Organization to Function, Cell Host & Microbe. 21 (2017) 433–442. https://doi.org/10.1016/j.chom.2017.03.010.

[3] J. Frampton, K.G. Murphy, G. Frost, E.S. Chambers, Short-chain fatty acids as potential regulators of skeletal muscle metabolism and function, Nat Metab. 2 (2020) 840–848. https://doi.org/10.1038/s42255-020-0188-7.

[4] J. Nguyen, D.M. Pepin, C. Tropini, Cause or effect? The spatial organization of pathogens and the gut microbiota in disease, Microbes and Infection. 23 (2021) 104815. https://doi.org/10.1016/j.micinf.2021.104815.

[5] D.J. Morrison, T. Preston, Formation of short chain fatty acids by the gut microbiota and their impact on human metabolism, Gut Microbes. 7 (2016) 189–200. https://doi.org/10.1080/19490976.2015.1134082.

[6] M.A.R. Vinolo, H.G. Rodrigues, R.T. Nachbar, R. Curi, Regulation of Inflammation by Short Chain Fatty Acids, Nutrients. 3 (2011) 858–876. https://doi.org/10.3390/nu3100858.

[7] B. Dalile, L. Van Oudenhove, B. Vervliet, K. Verbeke, The role of short-chain fatty acids in microbiota–gut–brain communication, Nat Rev Gastroenterol Hepatol. 16 (2019) 461–478. https://doi.org/10.1038/s41575-019-0157-3.

[8] E.E. Canfora, J.W. Jocken, E.E. Blaak, Short-chain fatty acids in control of body weight and insulin sensitivity, Nat Rev Endocrinol. 11 (2015) 577–591. https://doi.org/10.1038/nrendo.2015.128.

[9] S. Lucas, Y. Omata, J. Hofmann, M. Böttcher, A. Iljazovic, K. Sarter, O. Albrecht, O. Schulz, B. Krishnacoumar, G. Krönke, M. Herrmann, D. Mougiakakos, T. Strowig, G. Schett, M.M. Zaiss, Short-chain fatty acids regulate systemic bone mass and protect from pathological bone loss, Nat Commun. 9 (2018) 55. https://doi.org/10.1038/s41467-017-02490-4.

[10] S. Sanna, N.R. van Zuydam, A. Mahajan, A. Kurilshikov, A. Vich Vila, U. Võsa, Z. Mujagic, A.A.M. Masclee, D.M.A.E. Jonkers, M. Oosting, L.A.B. Joosten, M.G. Netea, L. Franke, A. Zhernakova, J. Fu, C. Wijmenga, M.I. McCarthy, Causal relationships among the gut microbiome, short-chain fatty acids and metabolic diseases, Nat Genet. 51 (2019) 600–605. https://doi.org/10.1038/s41588-019-0350-x.

[11] A.T. James, A.J.P. Martin, Gas-liquid partition chromatography: the separation and microestimation of volatile fatty acids from formic acid to dodecanoic acid, Biochemical Journal. 50 (1952) 679–690. https://doi.org/10.1042/bj0500679.

[12] D. Fiorini, M.C. Boarelli, R. Gabbianelli, R. Ballini, D. Pacetti, A quantitative headspace– solid-phase microextraction–gas chromatography–flame ionization detector method to analyze short chain free fatty acids in rat feces, Analytical Biochemistry. 508 (2016) 12–14. https://doi.org/10.1016/j.ab.2016.05.023.

[13] L. He, M.A.I. Prodhan, F. Yuan, X. Yin, P.K. Lorkiewicz, X. Wei, W. Feng, C. McClain, X. Zhang, Simultaneous quantification of straight-chain and branched-chain short chain fatty acids by gas chromatography mass spectrometry, Journal of Chromatography B. 1092 (2018) 359–367. https://doi.org/10.1016/j.jchromb.2018.06.028.

[14] M. Primec, D. Mičetić-Turk, T. Langerholc, Analysis of short-chain fatty acids in human feces: A scoping review, Analytical Biochemistry. 526 (2017) 9–21. https://doi.org/10.1016/j.ab.2017.03.007.

[15] G. Zhao, M. Nyman, J. Åke Jönsson, Rapid determination of short-chain fatty acids in colonic contents and faeces of humans and rats by acidified water-extraction and direct-injection gas chromatography, Biomed. Chromatogr. 20 (2006) 674–682. https://doi.org/10.1002/bmc.580.

[16] E.S. Lima, D.S.P. Abdalla, High-performance liquid chromatography of fatty acids in biological samples, Analytica Chimica Acta. 465 (2002) 81–91. https://doi.org/10.1016/S0003-2670(02)00206-4.

[17] N. Larsen, F.K. Vogensen, R.J. Gøbel, K.F. Michaelsen, S.D. Forssten, S.J. Lahtinen, M. Jakobsen, Effect of Lactobacillus salivarius Ls-33 on fecal microbiota in obese adolescents, Clinical Nutrition. 32 (2013) 935–940. https://doi.org/10.1016/j.clnu.2013.02.007.

[18] S. Sierra, F. Lara-Villoslada, L. Sempere, M. Olivares, J. Boza, J. Xaus, Intestinal and immunological effects of daily oral administration of Lactobacillus salivarius CECT5713 to healthy adults, Anaerobe. 16 (2010) 195–200. https://doi.org/10.1016/j.anaerobe.2010.02.001.

[19] L. Achour, S. Nancey, D. Moussata, I. Graber, B. Messing, B. Flourié, Faecal bacterial mass and energetic losses in healthy humans and patients with a short bowel syndrome, Eur J Clin Nutr. 61 (2007) 233–238. https://doi.org/10.1038/sj.ejcn.1602496.

[20] A.L. McOrist, G.C.J. Abell, C. Cooke, K. Nyland, Bacterial population dynamics and faecal short-chain fatty acid (SCFA) concentrations in healthy humans, Br J Nutr. 100 (2008) 138–146. https://doi.org/10.1017/S0007114507886351.

[21] Z. Chen, Y. Wu, R. Shrestha, Z. Gao, Y. Zhao, Y. Miura, A. Tamakoshi, H. Chiba, S.-P. Hui, Determination of total, free and esterified short-chain fatty acid in human serum by liquid chromatography-mass spectrometry, Ann Clin Biochem. 56 (2019) 190–197. https://doi.org/10.1177/0004563218801393.

[22] J. Han, K. Lin, C. Sequeira, C.H. Borchers, An isotope-labeled chemical derivatization method for the quantitation of short-chain fatty acids in human feces by liquid chromatography–tandem mass spectrometry, Analytica Chimica Acta. 854 (2015) 86–94. https://doi.org/10.1016/j.aca.2014.11.015.

[23] M. Okazaki, S. Matsukuma, R. Suto, K. Miyazaki, M. Hidaka, M. Matsuo, S. Noshima, N. Zempo, T. Asahara, K. Nomoto, Perioperative synbiotic therapy in elderly patients undergoing gastroenterological surgery: A prospective, randomized control trial, Nutrition. 29 (2013) 1224–1230. https://doi.org/10.1016/j.nut.2013.03.015.

[24] S. Ohigashi, K. Sudo, D. Kobayashi, T. Takahashi, K. Nomoto, H. Onodera, Significant Changes in the Intestinal Environment After Surgery in Patients with Colorectal Cancer, J Gastrointest Surg. 17 (2013) 1657–1664. https://doi.org/10.1007/s11605-013-2270-x.

[25] G. Le Gall, S.O. Noor, K. Ridgway, L. Scovell, C. Jamieson, I.T. Johnson, I.J. Colquhoun, E.K. Kemsley, A. Narbad, Metabolomics of Fecal Extracts Detects Altered Metabolic Activity of Gut Microbiota in Ulcerative Colitis and Irritable Bowel Syndrome, J. Proteome Res. 10 (2011) 4208–4218. https://doi.org/10.1021/pr2003598.

[26] J. Kong, J. Shi, Z. Chen, Y. Zeng, H. Chang, J. Ye, Q. Chu, Tributyl phosphate assisted hollowfiber liquid-phase microextraction of short-chain fatty acids in microbial degradation fluid using capillary electrophoresis-contactless coupled conductivity detection, Journal of Pharmaceutical and Biomedical Analysis. 154 (2018) 191–197. https://doi.org/10.1016/j.jpba.2018.02.058.

[27] A. Yavarinasab, M. Abedini, H. Tahmooressi, S. Janfaza, N. Tasnim, M. Hoorfar, Potentiodynamic Electrochemical Impedance Spectroscopy of Polyaniline-Modified Pencil Graphite Electrodes for Selective Detection of Biochemical Trace Elements, Polymers. 14 (2021) 31. https://doi.org/10.3390/polym14010031.

[28] G.P. Mane, S.N. Talapaneni, C. Anand, S. Varghese, H. Iwai, Q. Ji, K. Ariga, T. Mori, A. Vinu, Preparation of Highly Ordered Nitrogen-Containing Mesoporous Carbon from a Gelatin Biomolecule and its Excellent Sensing of Acetic Acid, Adv. Funct. Mater. 22 (2012) 3596–3604. https://doi.org/10.1002/adfm.201200207.

[29] L. Ma, S.Y. Ma, Z. Qiang, X.L. Xu, Q. Chen, H.M. Yang, H. Chen, Q. Ge, Q.Z. Zeng, B.Q. Wang, Preparation of Co-doped LaFeO 3 nanofibers with enhanced acetic acid sensing properties, Materials Letters. 200 (2017) 47–50. https://doi.org/10.1016/j.matlet.2017.04.096.

[30] L. He, C. Gao, L. Yang, K. Zhang, X. Chu, S. Liang, D. Zeng, Facile synthesis of MgGa2O4/graphene composites for room temperature acetic acid gas sensing, Sensors and Actuators B: Chemical. 306 (2020) 127453. https://doi.org/10.1016/j.snb.2019.127453.

[31] N.J. Pineau, F. Krumeich, A.T. Güntner, S.E. Pratsinis, Y-doped ZnO films for acetic acid sensing down to ppb at high humidity, Sensors and Actuators B: Chemical. 327 (2021) 128843. https://doi.org/10.1016/j.snb.2020.128843.

[32] Y.-C. Wang, Z.-S. Sun, S.-Z. Wang, S.-Y. Wang, S.-X. Cai, X.-Y. Huang, K. Li, Z.-T. Chi, S.-D. Pan, W.-F. Xie, Sub-ppm acetic acid gas sensor based on In2O3 nanofibers, J Mater Sci. 54 (2019) 14055–14063. https://doi.org/10.1007/s10853-019-03877-y.

[33] Y. Zhang, J. Liu, X. Chu, S. Liang, L. Kong, Preparation of g–C3N4–SnO2 composites for application as acetic acid sensor, Journal of Alloys and Compounds. 832 (2020) 153355. https://doi.org/10.1016/j.jallcom.2019.153355.

[34] C. Wang, S. Ma, A. Sun, R. Qin, F. Yang, X. Li, F. Li, X. Yang, Characterization of electrospun Pr-doped ZnO nanostructure for acetic acid sensor, Sensors and Actuators B: Chemical. 193 (2014) 326–333. https://doi.org/10.1016/j.snb.2013.11.058.

[35] Z. Su, X. Xu, Y. Cheng, Y. Tan, L. Xiao, D. Tang, H. Jiang, X. Qin, H. Wang, Chemical pre-reduction and electro-reduction guided preparation of a porous graphene bionanocomposite for indole-3-acetic acid detection, Nanoscale. 11 (2019) 962–967. https://doi.org/10.1039/C8NR06913A.

[36] G. Fusco, F. Gallo, C. Tortolini, P. Bollella, F. Ietto, A. De Mico, A. D’Annibale, R. Antiochia, G. Favero, F. Mazzei, AuNPs-functionalized PANABA-MWCNTs nanocomposite-based impedimetric immunosensor for 2,4-dichlorophenoxy acetic acid detection, Biosensors and Bioelectronics. 93 (2017) 52–56. https://doi.org/10.1016/j.bios.2016.10.016.

[37] C.-J. Fang, H.-C. You, Z.-L. Huang, C.-L. Hsu, C.-F. Tsai, Y.-T. Lin, Y.-M. Kao, S.-H. Tseng, D.-Y. Wang, N.-W. Su, Simultaneous Analysis of the Stable Carbon Isotope Ratios of Acetoin and Acetic Acid by GC-C-IRMS for Adulteration Detection in Brewed Rice Vinegar Products, J. Agric. Food Chem. 68 (2020) 14252–14260. https://doi.org/10.1021/acs.jafc.0c05674.

[38] S. Supharoek, K. Ponhong, W. Siriangkhawut, K. Grudpan, Employing natural reagents from turmeric and lime for acetic acid determination in vinegar sample, Journal of Food and Drug Analysis. 26 (2018) 583–590. https://doi.org/10.1016/j.jfda.2017.06.007.

[39] S. Han, W. Van Treuren, C.R. Fischer, B.D. Merrill, B.C. DeFelice, J.M. Sanchez, S.K. Higginbottom, L. Guthrie, L.A. Fall, D. Dodd, M.A. Fischbach, J.L. Sonnenburg, A metabolomics pipeline for the mechanistic interrogation of the gut microbiome, Nature. 595 (2021) 415–420. https://doi.org/10.1038/s41586-021-03707-9.

[40] J.C. De MAN, M. Rogosa, M.E. Sharpe, A MEDIUM FOR THE CULTIVATION OF LACTOBACILLI, Journal of Applied Bacteriology. 23 (1960) 130–135. https://doi.org/10.1111/j.1365-2672.1960.tb00188.x.

[41] J.S. Johnson, D.J. Spakowicz, B.-Y. Hong, L.M. Petersen, P. Demkowicz, L. Chen, S.R. Leopold, B.M. Hanson, H.O. Agresta, M. Gerstein, E. Sodergren, G.M. Weinstock, Evaluation of 16S rRNA gene sequencing for species and strain-level microbiome analysis, Nat Commun. 10 (2019) 5029. https://doi.org/10.1038/s41467-019-13036-1.

[42] R.E. Ley, J.K. Harris, J. Wilcox, J.R. Spear, S.R. Miller, B.M. Bebout, J.A. Maresca, D.A. Bryant, M.L. Sogin, N.R. Pace, Unexpected Diversity and Complexity of the Guerrero Negro Hypersaline Microbial Mat, Appl Environ Microbiol. 72 (2006) 3685–3695. https://doi.org/10.1128/AEM.72.5.3685-3695.2006.

[43] Thomas Madden, The BLAST Sequence Analysis Tool, National Center for Biotechnology Information (US), 2013. https://www.ncbi.nlm.nih.gov/books/NBK21097/.

[44] D. Lüdecke, ggeffects: Tidy Data Frames of Marginal Effects from Regression Models, JOSS. 3 (2018) 772. https://doi.org/10.21105/joss.00772.

[45] H. Wickham, ggplot2: Elegant Graphics for Data Analysis, 2nd ed. 2016, Springer International Publishing : Imprint: Springer, Cham, 2016. https://doi.org/10.1007/978-3-319-24277-4.

[46] P.V. Gaikwad, S.K. Sharma, K. Sudarshan, V. Kumar, A. Kshirsagar, P.K. Pujari, Molecular packing of polyvinyl alcohol in PVA-gold nanoparticles composites and its role on thermo-mechanical properties, Polym. Compos. 39 (2018) 1137–1143. https://doi.org/10.1002/pc.24042.

[47] M. Rau, A. Rehman, M. Dittrich, A.K. Groen, H.M. Hermanns, F. Seyfried, N. Beyersdorf, T. Dandekar, P. Rosenstiel, A. Geier, Fecal SCFAs and SCFA-producing bacteria in gut microbiome of human NAFLD as a putative link to systemic T-cell activation and advanced disease, United European Gastroenterol. j. 6 (2018) 1496–1507. https://doi.org/10.1177/2050640618804444.

[48] J. Tao, J. Duan, S. Jiang, J. Guo, Y. Qian, D. Qian, Simultaneous determination of six short-chain fatty acids in colonic contents of colitis mice after oral administration of polysaccharides from Chrysanthemum morifolium Ramat by gas chromatography with flame ionization detector, Journal of Chromatography B. 1029–1030 (2016) 88–94. https://doi.org/10.1016/j.jchromb.2016.07.002.

[49] L. Zhu, W. Zeng, Room-temperature gas sensing of ZnO-based gas sensor: A review, Sensors and Actuators A: Physical. 267 (2017) 242–261. https://doi.org/10.1016/j.sna.2017.10.021.

[50] D.-W. Kang, Z.E. Ilhan, N.G. Isern, D.W. Hoyt, D.P. Howsmon, M. Shaffer, C.A. Lozupone, J. Hahn, J.B. Adams, R. Krajmalnik-Brown, Differences in fecal microbial metabolites and microbiota of children with autism spectrum disorders, Anaerobe. 49 (2018) 121–131. https://doi.org/10.1016/j.anaerobe.2017.12.007.

[51] H.K. Seitz, U. Gärtner, G. Egerer, U.A. Simanowski, Ethanol metabolism in the gastrointestinal tract and its possible consequences, Alcohol Alcohol Suppl. 2 (1994) 157–162.

[52] A. Yavarinasab, S. Janfaza, H. Tahmooressi, M. Ghazi, N. Tasnim, M. Hoorfar, A selective polypyrrole-based sub-ppm impedimetric sensor for the detection of dissolved hydrogen sulfide and ammonia in a mixture, Journal of Hazardous Materials. 416 (2021) 125892. https://doi.org/10.1016/j.jhazmat.2021.125892.

[53] S. Brosel-Oliu, N. Abramova, N. Uria, A. Bratov, Impedimetric transducers based on interdigitated electrode arrays for bacterial detection – A review, Analytica Chimica Acta. 1088 (2019) 1–19. https://doi.org/10.1016/j.aca.2019.09.026.

[54] J.-B. Jorcin, M.E. Orazem, N. Pébère, B. Tribollet, CPE analysis by local electrochemical impedance spectroscopy, Electrochimica Acta. 51 (2006) 1473–1479. https://doi.org/10.1016/j.electacta.2005.02.128.

[55] H. Ma, J. Li, X. Cheng, A. Nathan, Heterogeneously integrated impedance measuring system with disposable thin-film electrodes, Sensors and Actuators B: Chemical. 211 (2015) 77–82. https://doi.org/10.1016/j.snb.2015.01.044.

[56] X.B. Li, Q.Q. Zhang, S.Y. Ma, G.X. Wan, F.M. Li, X.L. Xu, Microstructure optimization and gas sensing improvement of ZnO spherical structure through yttrium doping, Sensors and Actuators B: Chemical. 195 (2014) 526–533. https://doi.org/10.1016/j.snb.2014.01.087.

[57] G. Li, Y. Su, Y.-Y. Li, Y.-X. Li, Z. Guo, X.-J. Huang, J.-H. Liu, Size-tunable Ag nanoparticles sensitized porous ZnO nanobelts: controllably partial cation-exchange synthesis and selective sensing toward acetic acid, Nanotechnology. 29 (2018) 445501. https://doi.org/10.1088/1361-6528/aada6e.

[58] V. Khorramshahi, J. Karamdel, R. Yousefi, Acetic acid sensing of Mg-doped ZnO thin films fabricated by the sol–gel method, J Mater Sci: Mater Electron. 29 (2018) 14679–14688. https://doi.org/10.1007/s10854-018-9604-0.

[59] W.X. Jin, S.Y. Ma, Z.Z. Tie, W.Q. Li, J. Luo, L. Cheng, X.L. Xu, T.T. Wang, X.H. Jiang, Y.Z. Mao, Synthesis of hierarchical SnO2 nanoflowers with enhanced acetic acid gas sensing properties, Applied Surface Science. 353 (2015) 71–78. https://doi.org/10.1016/j.apsusc.2015.06.089.

[60] L. Cheng, S.Y. Ma, T.T. Wang, J. Luo, X.B. Li, W.Q. Li, Y.Z. Mao, D.J. Gz, Highly sensitive acetic acid gas sensor based on coral-like and Y-doped SnO2 nanoparticles prepared by electrospinning, Materials Letters. 137 (2014) 265–268. https://doi.org/10.1016/j.matlet.2014.09.040.

[61] M. Turemis, D. Zappi, M.T. Giardi, G. Basile, A. Ramanaviciene, A. Kapralovs, A. Ramanavicius, R. Viter, ZnO/polyaniline composite based photoluminescence sensor for the determination of acetic acid vapor, Talanta. 211 (2020) 120658. https://doi.org/10.1016/j.talanta.2019.120658.

[62] J. Chen, A.A. Jackson, V.M. Rotello, S.R. Nugen, Colorimetric Detection of Escherichia coli Based on the Enzyme-Induced Metallization of Gold Nanorods, Small. 12 (2016) 2469–2475. https://doi.org/10.1002/smll.201503682.

[63] A. Yavarinasab, S. Janfaza, N. Tasnim, H. Tahmooressi, A. Dalili, M. Hoorfar, Graphene/poly (methyl methacrylate) electrochemical impedance-transduced chemiresistor for detection of volatile organic compounds in aqueous medium, Analytica Chimica Acta. 1109 (2020) 27–36. https://doi.org/10.1016/j.aca.2020.02.065.

[64] G. D’Argenio, G. Mazzacca, Short-Chain Fatty Acid in the Human Colon, in: V. Zappia, F. Della Ragione, A. Barbarisi, G.L. Russo, R.D. Iacovo (Eds.), Advances in Nutrition and Cancer 2, Springer US, Boston, MA, 1999: pp. 149–158. https://doi.org/10.1007/978-1-4757-3230-6_13.

[65] J.G. LeBlanc, F. Chain, R. Martín, L.G. Bermúdez-Humarán, S. Courau, P. Langella, Beneficial effects on host energy metabolism of short-chain fatty acids and vitamins produced by commensal and probiotic bacteria, Microb Cell Fact. 16 (2017) 79. https://doi.org/10.1186/s12934-017-0691-z.

[66] A. Nakkarach, H.L. Foo, A.A.-L. Song, N.E.A. Mutalib, S. Nitisinprasert, U. Withayagiat, Anticancer and anti-inflammatory effects elicited by short chain fatty acids produced by Escherichia coli isolated from healthy human gut microbiota, Microb Cell Fact. 20 (2021) 36. https://doi.org/10.1186/s12934-020-01477-z.

[67] A. Nakkarach, H.L. Foo, A.A.-L. Song, S. Nitisinprasert, U. Withayagiat, Promising discovery of beneficial Escherichia coli in the human gut, 3 Biotech. 10 (2020) 296. https://doi.org/10.1007/s13205-020-02289-z.

[68] K. Sivieri, M.L.V. Morales, M.A.T. Adorno, I.K. Sakamoto, S.M.I. Saad, E.A. Rossi, Lactobacillus acidophilus CRL 1014 improved “gut health” in the SHIME®reactor, BMC Gastroenterol. 13 (2013) 100. https://doi.org/10.1186/1471-230X-13-100.

[69] M. Müller, M.A.G. Hernández, G.H. Goossens, D. Reijnders, J.J. Holst, J.W.E. Jocken, H. van Eijk, E.E. Canfora, E.E. Blaak, Circulating but not faecal short-chain fatty acids are related to insulin sensitivity, lipolysis and GLP-1 concentrations in humans, Sci Rep. 9 (2019) 12515. https://doi.org/10.1038/s41598-019-48775-0.

