## Supplementary figures and images for "An impedance-based chemiresistor for the real-time detection of gut microbiota-generated short-chain fatty acids"

### Graphical abstract

#### Graphical abstract

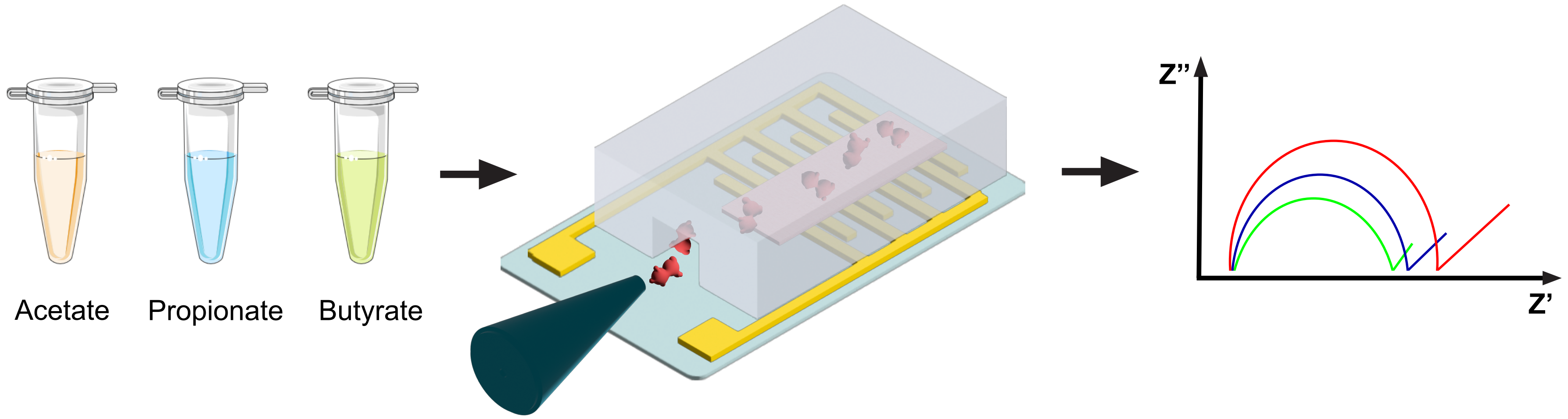

### Supplemental Figure 1

#### Figure S1

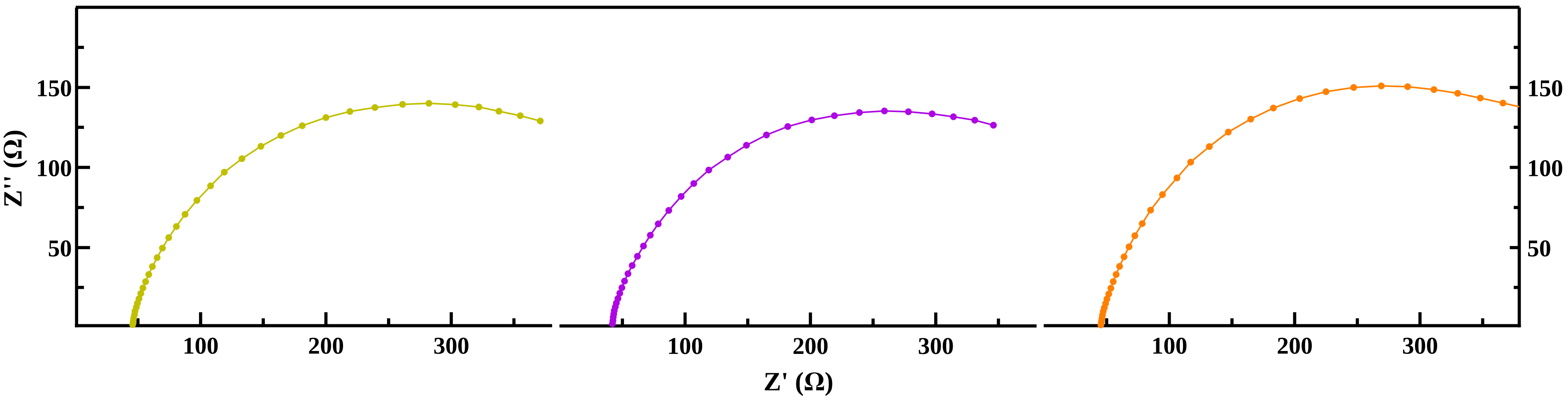

### Supplemental Figure 2

#### Figure S2

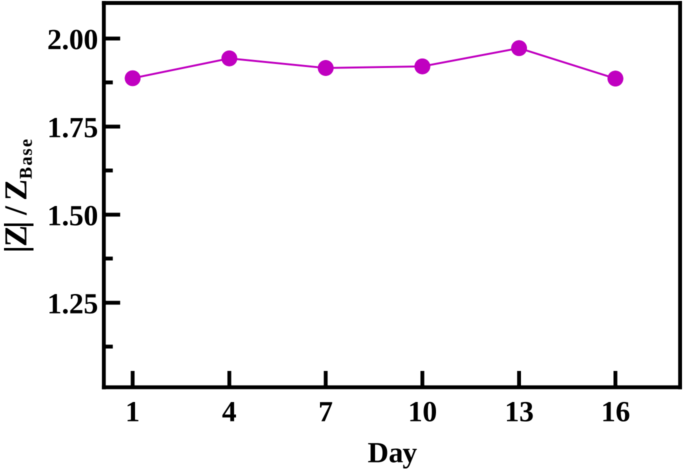

### Supplemental Figure 3

#### Figure S3

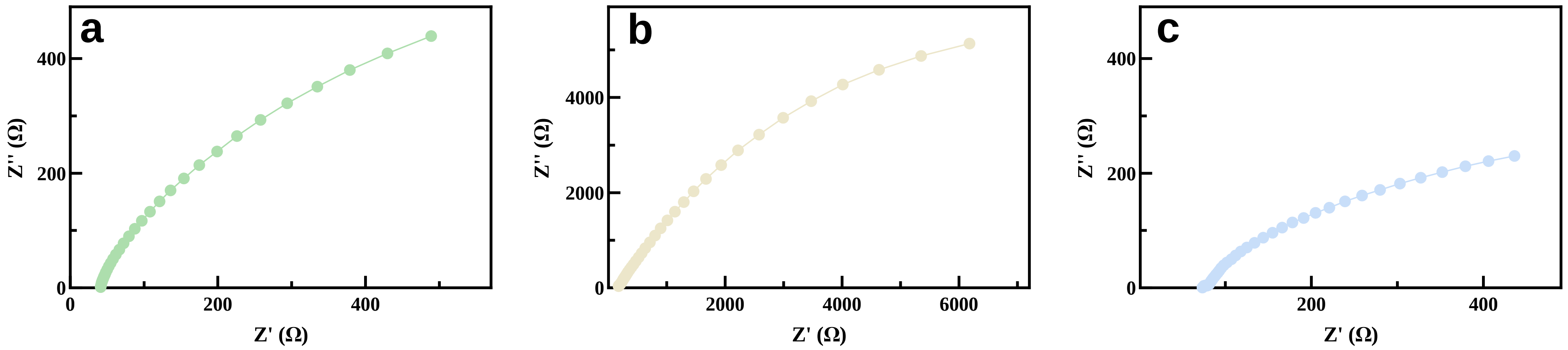

### Supplemental Figure 4

#### Figure S4

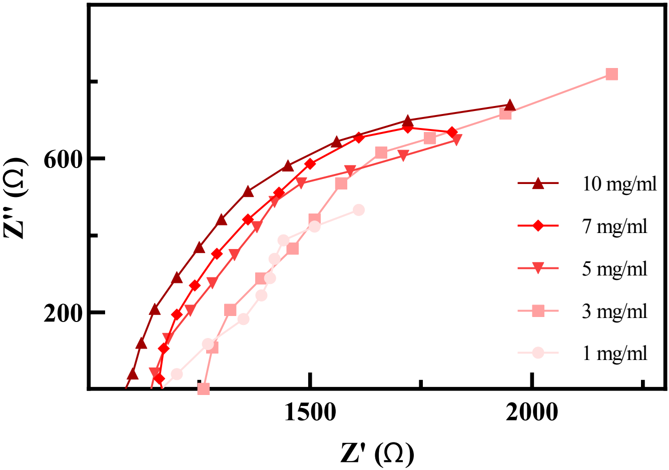

### Supplemental Figure 5

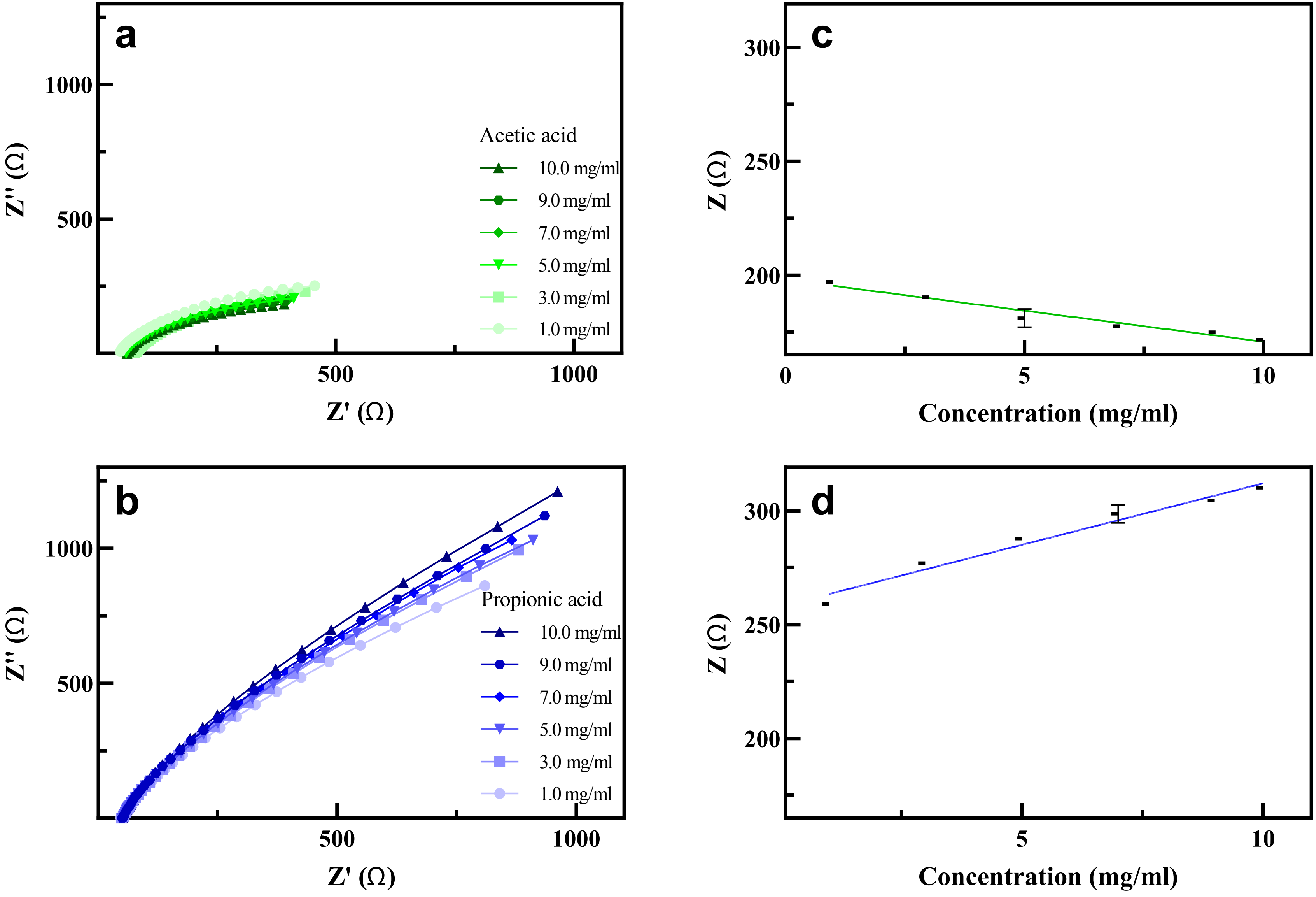

### Supplemental Figure 6

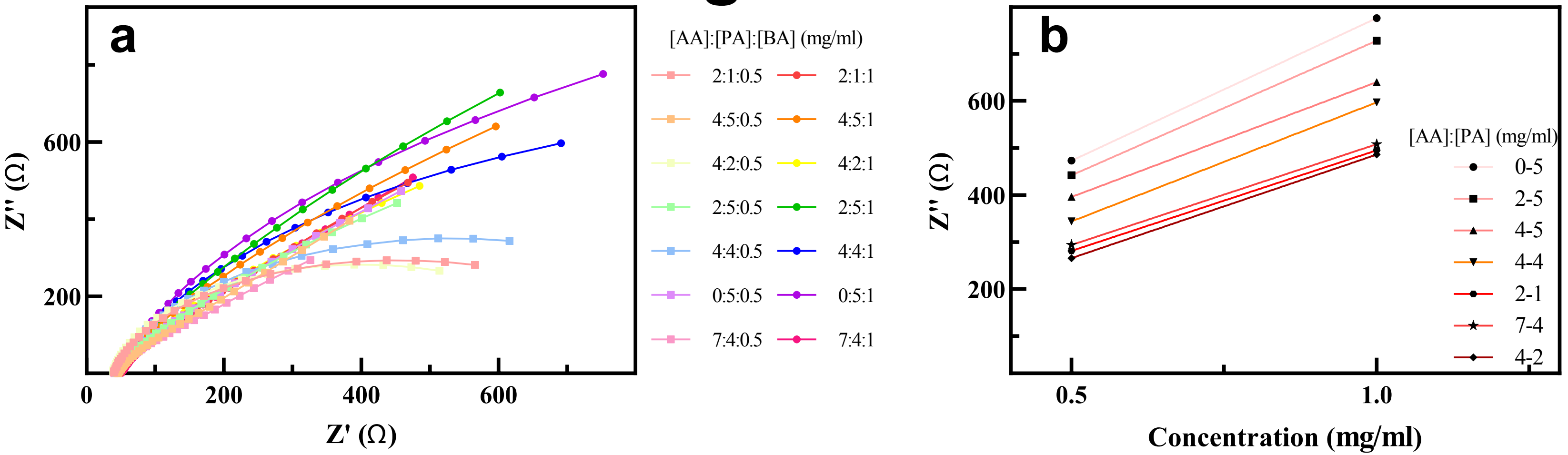

### Supplemental Figure 7

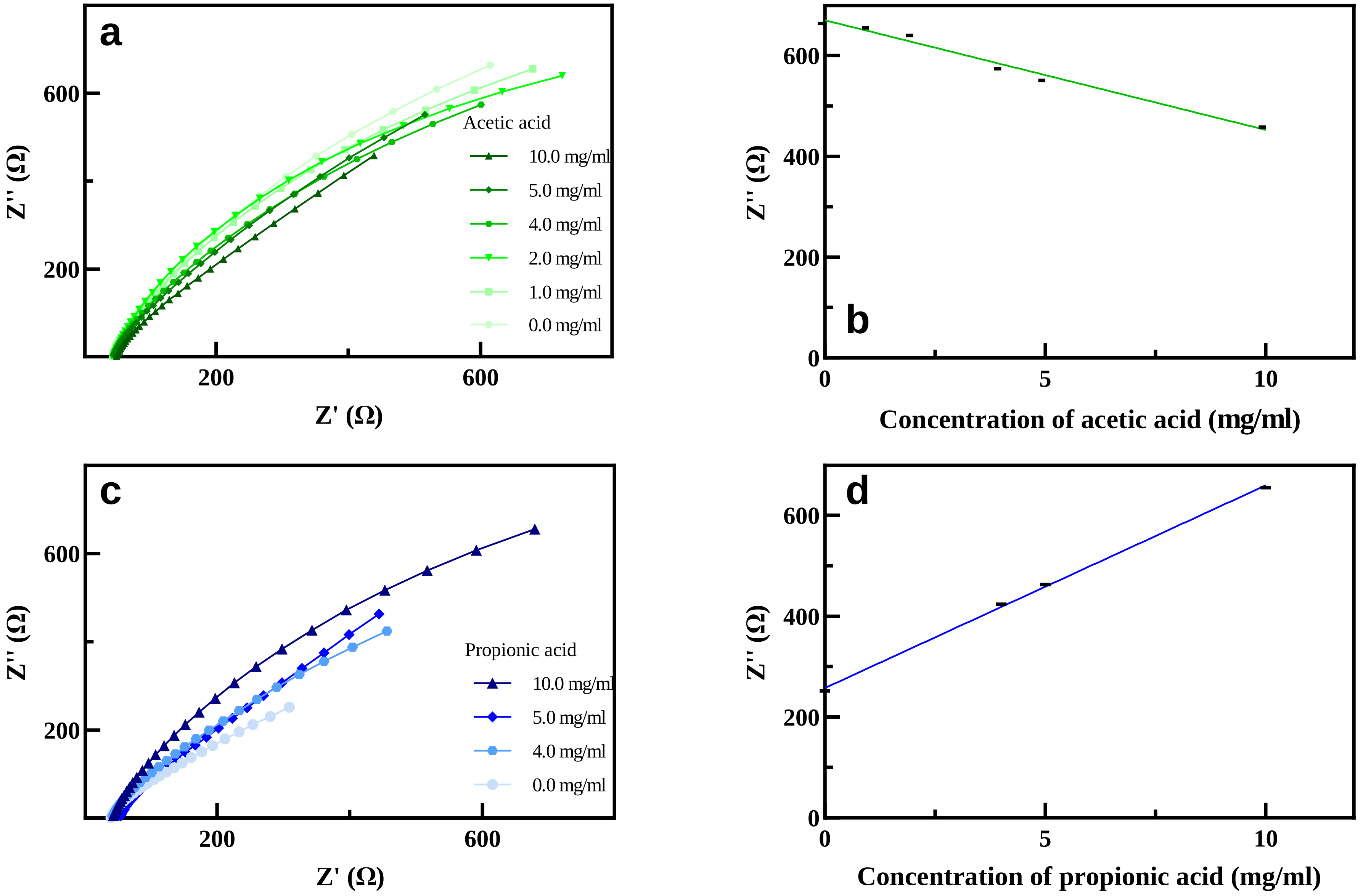
